# Cell-specific regulatory circuits connect genetic variation to disease susceptibility

**DOI:** 10.64898/2026.06.01.729215

**Authors:** Roy Oelen, Maryna Korshevniuk, Jelmer Niewold, Dan Kaptijn, Martijn van der Werff, Marc Jan Bonder, sc-eQTLGen Consortium, Lude Franke, Monique G.P. van der Wijst

## Abstract

Genome-wide association studies have identified thousands of variants associated with immune-related diseases, yet most lie in non-coding regions, complicating mechanistic interpretation. Regulatory quantitative trait loci (QTLs), such as expression QTLs (eQTLs) and chromatin accessibility QTLs (caQTLs), offer a powerful framework for prioritization and interpretation of these disease-associated genetic variants. When analyzed together, they offer deeper insights into the regulatory architecture underlying disease.

We generated same-cell, single-cell multi-omics data, integrating transcriptomic and chromatin accessibility information, from 563,100 matched peripheral blood mononuclear cells collected from 264 individuals, either unstimulated or stimulated for 24h with *C. albicans* (CA). Across six major immune cell types, we mapped both *cis*-eQTLs and -caQTLs, identifying 1,571 eGenes and 28,862 caPeaks, with 41% and 11% showing a stimulation-dependent effect. Finally, to dissect the regulatory mechanisms underlying these QTL effects, we applied two complementary strategies: 1. overlapping caQTLs with eQTLs; 2. applying SCENIC+ to identify regulatory triplets containing a transcription factor, the chromatin region it may bind to and the candidate target genes it thereby may regulate.

With the first approach, we identified 1,861 dual-acting QTLs. These dual-QTLs showed 1.9-fold stronger enrichment for immune-related disease associations than single-modality QTLs, highlighting their relevance for disease interpretation. With the second approach, we found 62,932 regulatory triplets, of which 1.7% were under genetic control. By then leveraging the SCENIC+-derived TF activity measurements we could study how genetic variants can rewire TF control of gene expression, ultimately shaping inter-individual variation in disease risk.

Together, our network-based approach offers new insights into the cellular contexts and gene programs perturbed in disease, providing a foundation for prioritizing therapeutic targets and informing strategies for disease prevention.

## Introduction

Many diseases are driven by complex biological mechanisms involving the interplay between genetic and environmental risk factors. Genome-wide association studies (GWAS) have become a cornerstone in unraveling this complexity by identifying statistical associations between common genetic variants—primarily single nucleotide polymorphisms (SNPs)—and a wide range of traits or diseases (Uffelmann et al^1^). To date, over 200,000 genetic variants have been linked to more than 3,000 human traits, reflecting the growing depth of our genetic understanding^2^. However, this rapid expansion has underscored the critical need for functional interpretation and mechanistic validation of disease-associated variants^3^.

Variants within protein-coding regions are relatively straightforward to interpret, as they may directly alter the amino acid sequence, thereby potentially disrupting protein structure or function. In contrast, non-coding variants—though far more abundant—are more challenging to decipher. GWAS findings indicate that while coding variants are modestly enriched among fine-mapped loci, approximately 95% of the high-confidence SNPs lie within non-coding or flanking regions^4,5^. This striking pattern points to gene regulation as a central mechanism in disease susceptibility.

A key strategy to interpret the functional consequences of GWAS loci is through quantitative trait locus (QTL) mapping, which links genetic variants to changes in molecular phenotypes such as chromatin accessibility (caQTL), gene expression (eQTL), or protein abundance (pQTL). In the last decade, numerous large-scale research projects have been conducted to identify these QTLs at scale and across diverse ethnicities and tissues, such as eQTLGen^6^, GTEx^7^, and BLUEPRINT^8^ (<u>EBI eQTL</u> <u>Studies</u>). Until now, most of these efforts have focused on eQTLs, and used bulk omics data for mapping. However, more recently, also single-cell RNA-sequencing (scRNA-seq) data has been successfully applied, predominantly for eQTL mapping (Wijst 2018^9^, Oelen 2022^10^, Yazar et al., 2022^11^). Together, these analyses have revealed that many of these QTL effects are tissue-, cell-type-or even context-specific^7,10,12^, and therefore, relying on averaged bulk-derived QTLs may obscure or attenuate true regulatory signals^11,13^.

Importantly, previous analyses have highlighted that relying on a single omics layer alone, might not be sufficient to fully explain genetic disease susceptibility^12,14^. For example, the iPSCORE multiomic QTL study found that while eQTLs explained 43% of GWAS signals, integrating caQTLs increased locus annotation by 2.3-fold. These caQTLs likely reflect regulatory effects that are context-specific—where chromatin accessibility is already established (primed), but corresponding eQTLs only manifest in certain cell types or conditions^15^. These findings highlight the importance of considering multiple omics layers at once when interpreting genetic risk variants.

Only recently large-scale single-cell multiomics technologies have become available that jointly measure chromatin accessibility and gene expression in the same cell. These now enable us for the first time to dissect at scale how genetic variants impact gene regulation, while overcoming the risk of mitigating or misleading signals from averaging across heterogeneous cell populations.

In this study, we aim to better understand how genetic variants impact gene expression and its regulation. To achieve this, we performed same-cell, single-cell multiomics (scMO) profiling in peripheral blood mononuclear cells (PBMCs) from 264 donors. We then conducted QTL mapping across both molecular layers, enabling the detection of 1,571 independent expression QTL genes (eGenes) and 28,862 independent chromatin accessibility QTL peaks (caPeaks) across the 6 major PBMC cell types. Finally, we identified 62,932 regulatory triplets (transcription factor (TF), cis-regulatory element (CRE) and target gene), of which 17.7% was found to be under genetic control. Together, these network-level insights enhance our ability to interpret disease-related genetic variation, thereby advancing target discovery and contributing to more effective prevention strategies.

## Results

### Same-cell single-cell multiomic analyses in immune cells upon pathogen stimulation

Here we present a comprehensive multiomics analysis combining same-cell transcriptomic and epigenomic profiling of PBMCs from 264 healthy individuals from the northern Netherlands (age: 25-85 years, 59% female) (**Fig. 1a**). In total, 315 PBMC samples were analyzed, including samples either left untreated (128 individuals) or stimulated for 24h with *C. albicans* (24h CA) (187 individuals) (**Fig. 1b**). For this, we first pooled eight individuals in one sample pool, while aiming for a balanced composition of sex and condition. Nuclei were extracted from each pool and loaded in two separate lanes of a 10x Genomics Chromium Next GEM Single Cell Multiome ATAC + Gene Expression chip, targeting 2,500 PBMCs per sample. In parallel, we extracted DNA from these PBMCs and genotyped all individuals using the Illumina GSA-MD v3 array. After imputation this resulted in about 7 million genetic variants that were tested for genetic associations at a minor allele frequency (MAF) of ≥ 0.05.

**Figure 1.**
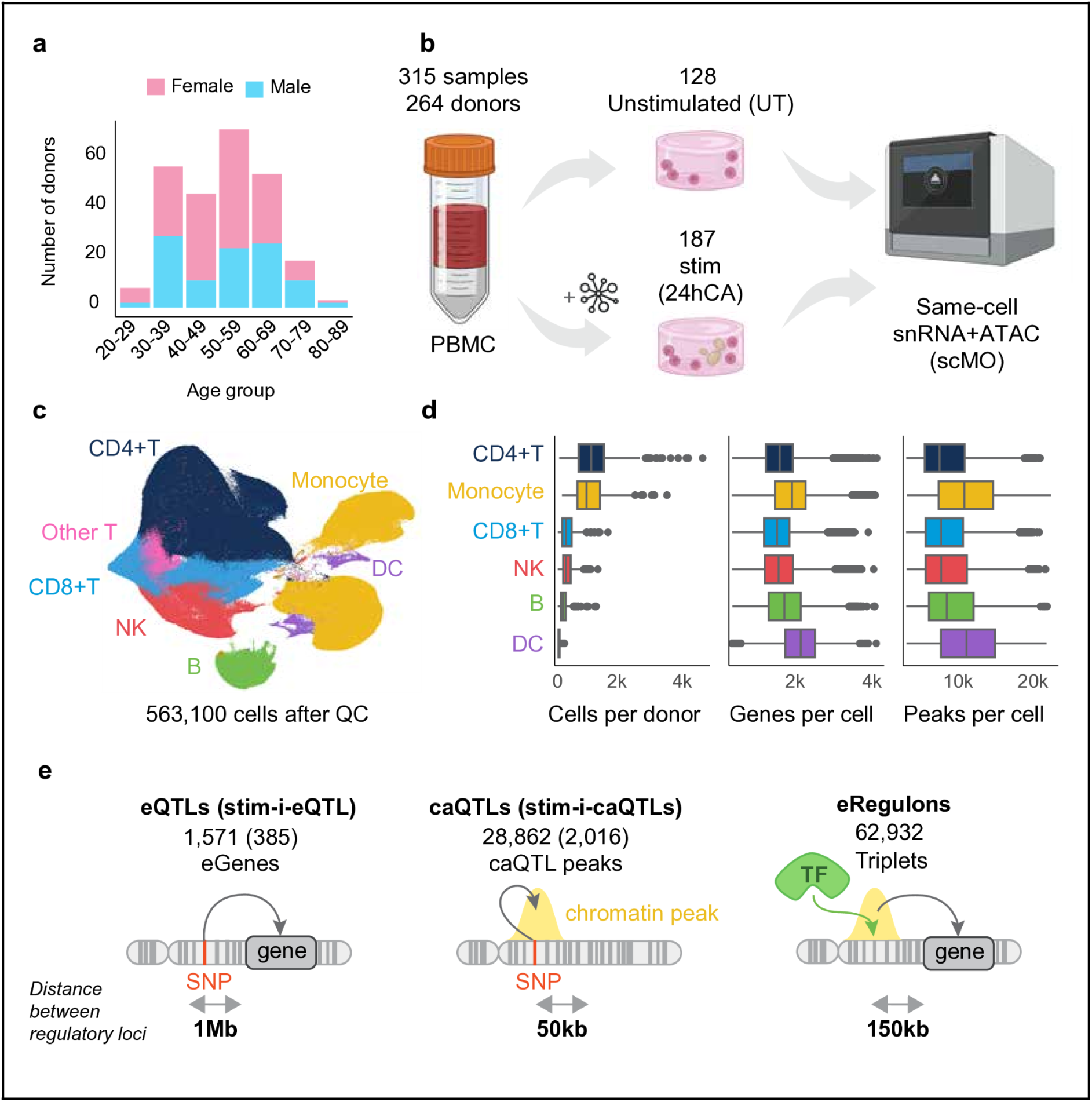
Experimental design and analytical workflow. **a, b.** Same-cell, single-cell multiome (scMO) data was generated from PBMCs from 264 healthy donors (25-85 years, 59% female): 128 unstimulated and 187 24h C. albicans (CA)-stimulated samples. **c.** Six major PBMC cell types were annotated with the 10x Multiome PBMC v1.0 reference using Azimuth. **d.** For each cell type the distributions are shown of the number of cells per donor, number of genes and peaks per cell. **e.** Downstream analyses conducted in this study: genetic variants were linked to expression (eQTL and stimulation-interaction-eQTL mapping) and chromatin accessibility (caQTL and stimulation-interaction-caQTL mapping), and this information was overlapped with identified regulatory triplets eRegulons (transcription factor (TF) - cis-regulatory element (CRE) - target gene).

We processed each modality using Cell Ranger ARC to align reads and generate initial feature-barcode matrices for gene expression and chromatin accessibility. We then removed ambient RNA contamination using CellBender^16^, assigned cells to donors by correlating donor genotypes to scRNA-seq-derived genotypes using SoupOrCell^17^, and applied quality control (QC) on the individual omics layers (**Suppl. Table 1**). Subsequently, cells were annotated using reference-based annotation.

For this, we first employed a weighted nearest-neighbour (WNN) approach to jointly embed scRNA-seq and scATAC-seq modalities^18^, and then annotated each cell using Azimuth with the 10x Multiome PBMC v1.0 reference. This yielded six major immune cell types (B, CD4+ T, CD8+ T, natural killer (NK), dendritic cell (DC) and monocyte) (**Fig. 1c**), of which CD4+ T cells and monocytes were the most abundant and DCs the least abundant (**Fig. 1d**). These cell type annotations were confirmed by assessing canonical marker gene expression of the assigned cell types (**Suppl. Fig. 1**). Altogether, we retained 563,100 high-quality cells from 264 individuals for which genetic, transcriptomic and epigenomic information was available. This data was used for downstream analyses (**Fig. 1e**).

### eQTLs and stim-i-eQTLs are mostly shared among cells from the same lineage

To identify genetic variants that affect gene expression levels, we conducted eQTL mapping in each of the six major PBMC cell types using the LIMIX-QTL pipeline^19^. This pipeline uses a mixed-effects model to associate variants to gene expression, while correcting for stimulation status and the first 20 principal components (PCs) of the pseudobulked gene expression as fixed effects, and donor and experimental batch as random effects. To then identify the independent credible sets likely harboring the causal variants that impact gene expression levels, we conducted fine-mapping using SuSiE^20^. Overall, this resulted in the identification of 1,571 unique eGenes, and an additional 178 non-primary eQTLs (range: 1-4 per eGene) (**Fig. 2a, Suppl. Table 2**). The number of eQTLs that were detected were closely related to the average number of cells per cell type and donor, with most eQTL effects identified in the CD4+ T cells, and least in the DCs (**Fig. 1d**, **2a**). Of the eGenes discovered, 75.1% were identified in only a single cell type, suggestive of apparent cell type-specificity. Partially, this cell type-specificity can be explained by the cell type-specific expression of the gene itself (**Suppl. Fig. 2**). On top of that, genes can be expressed among multiple cell types, but without sharing an eQTL effect on that gene. However, for this latter case we observed that effects are generally shared across immune cell types, as the pairwise comparison between eQTL effect sizes across immune cell types using the r*_b_*metric^21^ was generally high (r*_b_* 0.61-0.95), especially for cells that share the same lineage (**Fig. 2b**). These observations suggest that the observed cell type-specificity for the eQTLs may be partially driven by statistical power rather than true biological exclusivity. When we compared our eQTLs with the largest sc-eQTL PBMC study (OneK1K^11^, 982 individuals: cell type-matched, r*_b_* = 0.78-0.93) and the largest bulk-level whole blood dataset (eQTLGen consortium^6^ 31,682 individuals: r*_b_* = 0.47-0.57), again we observed high overlap in eQTL effect sizes, especially in matching cell types (**Fig. 2c, 2d, Suppl. Fig. 3a, 3b**). This high concordance across datasets suggests that our eQTL findings are robust.

**Figure 2.**
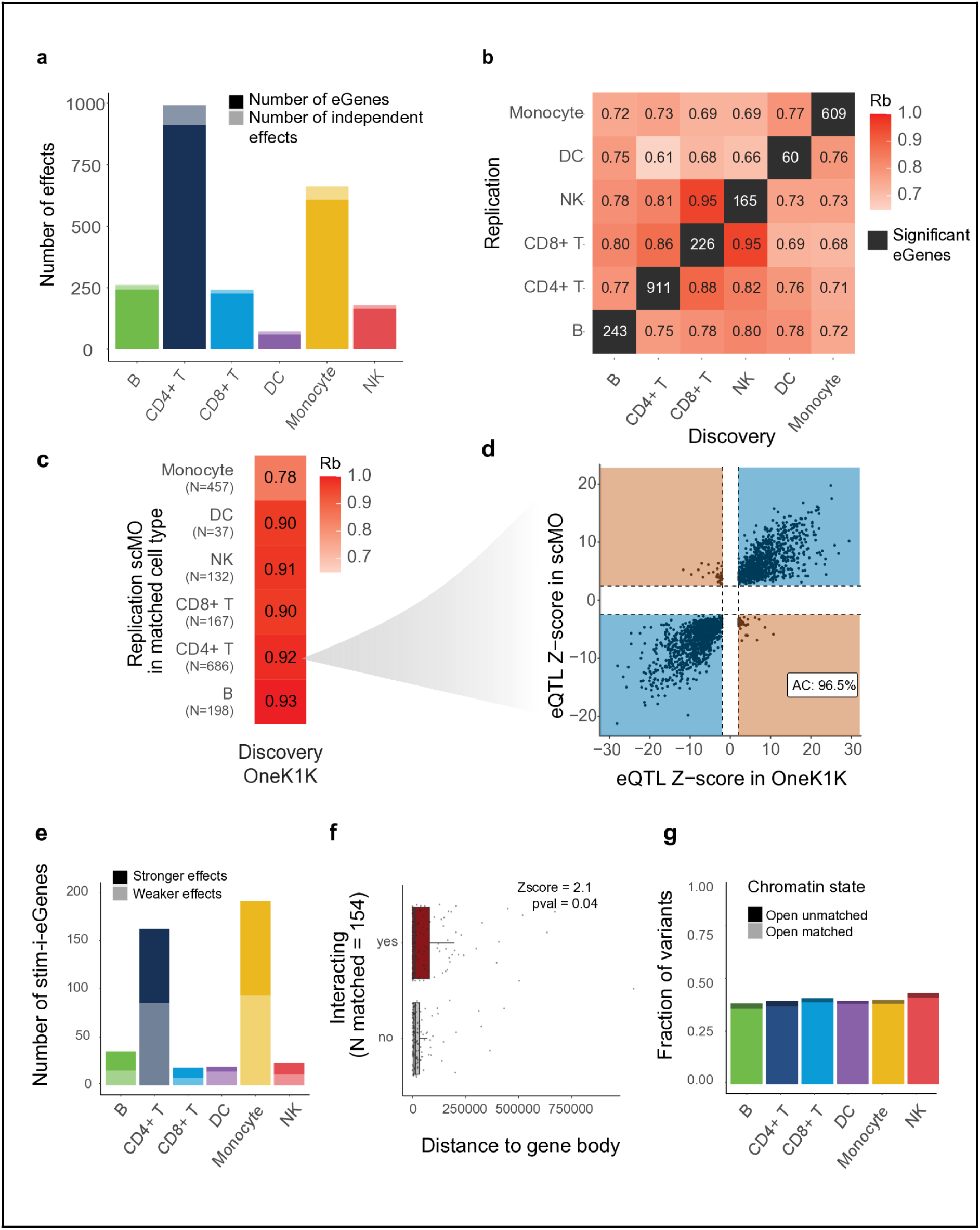
Expression QTLs and their interaction with 24h C. albicans stimulation. **a.** Number of eGenes and non-primary eQTLs identified per cell type. **b.** Heatmap showing the pairwise comparison of the eQTL effect sizes of the shared expressed genes as measured using the r_b_. **c.** Cell-type-matched concordance of the lead eQTLs identified in OneK1K^23^. **d.** Scatter plot showing the eQTL Z-score concordance of the eQTLs that were identified in OneK1K CD4+ T cells, and could be significantly replicated in our scMO CD4+ T cells. **e.** Number of stimulation-dependent interaction-eGenes identified per cell type, colored by whether the stimulation strengthened (dark tone) or weakened (light tone) the eQTL effect. **f.** Boxplots showing the distance between the lead eSNP and gene body, split by whether the SNP interacted with stimulation or not. SNPs are matched based on MAF, gene body size and eQTL effect size. **g.** Fraction of lead eSNPs in which the chromatin is open in the context in which the eQTL acts (matched) or only in unmatched context (unmatched), split by cell type.

Subsequently, for the eQTLs that were found in unstimulated or 24h CA-stimulated cells, we performed an interaction-eQTL analysis using the stimulation-status as interaction term, while correcting for the same variables as the baseline eQTL mapping. Overall, 41% of the eGenes also showed an interaction effect with stimulation status (stim-i-eGenes) (**Fig. 2e**), with about equal numbers of weakening and strengthening effects. Notably, the fraction of stim-i-eGenes seemed relatively higher for the myeloid compared to the lymphoid cell types, aligned with their higher responsiveness to the 24h CA stimulation (**Suppl. Fig. 4a**, **Suppl. Table 3**), consistent with our previous observation^10^. There was no strong bias between de DE and stim-i-eQTL direction (**Suppl. Fig. 4b**). This strong correlation between being an stim-i-eGene and DE gene, supports the robustness of our stim-i-eQTLs.

Finally, comparing the characteristics of the eQTLs with the stim-i-eQTLs, we noticed that on average the stim-i-eQTLs were located more distal from the gene body (p = 0.04, z = 2.1) (**Fig. 2f**). These findings reinforce the idea that distal regulatory elements such as enhancers play a key regulatory role in shaping context-dependent gene expression^22^.

### Open chromatin data can be used to provide additional interpretation of genetic variants

As open chromatin data can be used to further prioritize the likely causal variant and provide interpretation of the underlying mechanism of action that underlies an eQTL effect, we then explored the chromatin layer separately and together with the expression layer. For this, we first quantified the chromatin accessibility in each cell by overlapping our ATAC reads with peaks derived from a human reference set of consensus open chromatin regions (cPeaks)^24^. Taking advantage of such a reference, we can detect potential open chromatin regions with higher sensitivity and these peaks can more easily be compared and integrated with future datasets that use the same reference. Across all cells, we detected approximately 1.2 million peaks present in at least one of the six major cell types (out of the 1.68 million assessed cPeaks) (**Suppl. Note 1**). To distinguish called peaks from background noise, we restricted our further analyses to regions with sufficient coverage in that cell type or condition.

We found that selecting peaks that were detected in at least 0.1% of the cells of a particular major cell type, strikes the best balance between peak sensitivity and biological (i.e. driven by lineage-specificity, remain housekeeping peaks) over technical signal (i.e. driven by differences in statistical power) (**Suppl. Note 1**). This resulted in the identification of 668,059 high-confidence peaks across all cell types and conditions, of which 20.0% were more and 18.4% were less accessible after stimulation (**Suppl. Table 4**).

Chromatin often needs to open before expression can take place. Therefore, we checked whether our eQTL variants were located in open chromatin in the cell type in which the eQTL effect takes place. Overall, this was the case for 42% of the lead eQTL variants (**Fig. 2g**). This supports the hypothesis that many of the regulatory eQTL variants are located in open chromatin to exert their function.

Alternatively, genetic variants may directly regulate chromatin openness. To investigate this, we performed caQTL mapping across the six major PBMC cell types using the LIMIX-QTL pipeline^25^. This pipeline uses a mixed-effects model to associate variants to chromatin accessibility, while correcting for stimulation status and the first 20 PCs of the pseudobulked chromatin accessibility as fixed effects, and donor and experimental batch as random effects. We then identified the independent credible sets likely harboring the causal variants that impact chromatin accessibility levels using SuSiE^20^. Overall, this led to the identification of 28,862 unique caPeaks, of which 3.3% had an additional non-primary caQTLs (range: 1-3 per caPeak) (**Fig. 3a**, **Suppl. Table 5**). Of the caPeaks discovered, 73.7% were identified in only a single cell type, suggestive of apparent cell type-specificity. Partially, this cell type-specificity can be explained by the cell type-specific chromatin accessibility of the peak itself (**Suppl. Fig. 5**). On top of that, peaks can be accessible among multiple cell types, but without sharing a caQTL effect on that peak. In pairwise comparisons of caQTL effect sizes across immune cell types, we found that the overlap was generally high among cells from the same lineage (r*_b_* = .85-.92) (**Fig. 3b**), and that caQTLs showed slightly stronger lineage-specificity than eQTLs (**Fig. 2b**). This aligns with the findings of Kanai et al., who reported that the majority of eQTLs (73%) and caQTLs (77%) are (likely) shared among lineages, and that much of the apparent cell type-specificity can be attributed to differences in statistical power^26^. Next, to validate our caQTLs, we compared them with an external bulk-level lymphoblastoid cell line (LCL) caQTL dataset^27^ (**Fig. 3c**). Reassuringly, we observed the highest concordance in the B cells (r*_b_* = 0.61, 97.2% allelic concordance) —the most biologically similar cell type to LCLs (**Fig. 3d**), underscoring the robustness of our results.

**Figure 3.**
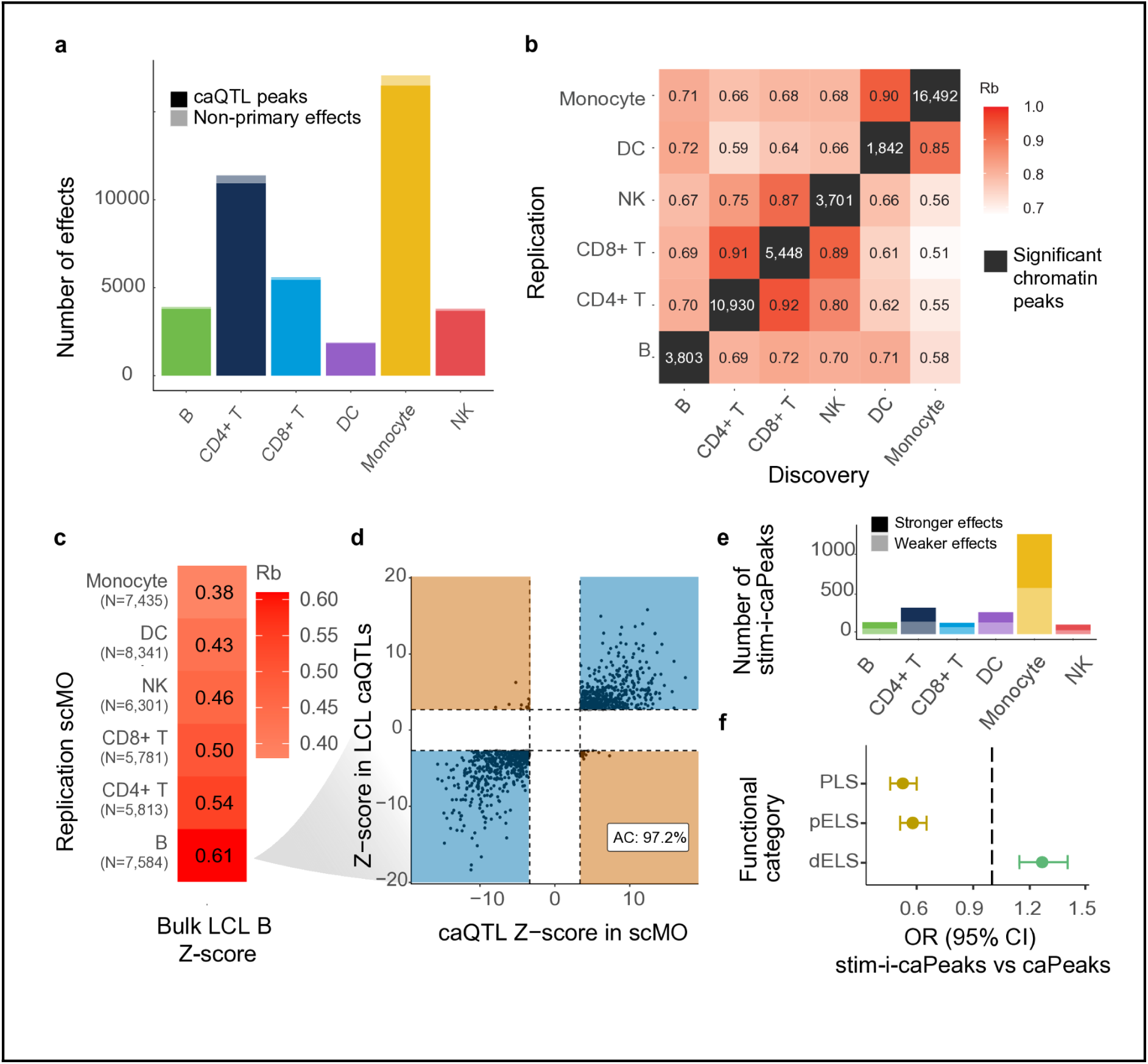
Chromatin accessibility QTLs and their interaction with 24h C. albicans stimulation. **a.** Number of caPeaks and non-primary caQTLs identified per cell type. **b.** Heatmap showing the pairwise comparison of the caQTL effect sizes of the shared accessible peaks as measured using the r_b_. **c.** Cell-type-matched concordance of the lead caQTLs identified in bulk LCL B data. **d.** Scatter plot showing the caQTL Z-score concordance of the caQTLs that were identified in bulk LCL B data, and significant in our scMO B cells. **e.** Number of stimulation-dependent interaction-caPeaks identified per cell type, colored by whether the stimulation strengthened (dark tone) or weakened (light tone) the caQTL effect. **f.** Enrichment of the stimulation-dependent interaction-caPeaks vs baseline caPeaks in distal (dELS: distal enhancer-like sequence) and proximal (pELS: proximal enhancer-like sequence, PLS: promoter-like sequence) regulatory elements.

Subsequently, we used the caQTLs that were found in unstimulated or 24h CA-stimulated cells for an interaction-caQTL analysis. For this, we used the stimulation-status as an interaction term, while adjusting for the same variables as the baseline caQTL mapping. Overall, 11.4% of the caPeaks also showed an interaction effect with stimulation status (stim-i-caPeaks), about equally likely strengthening or weakening the caQTL effect (**Fig. 3e**). Comparing the characteristics of the caPeaks with the stim-i-caPeaks, the stim-i-caPeaks were enriched for distal regulatory elements (dELS: distal enhancer-like sequences) and the caPeaks for proximal regulatory elements (pELS: proximal enhancer-like, PLS: promoter-like sequences)^28^ (**Fig. 3f**). This again supports a more complex regulation of context-specific effects, likely involving distal regulatory elements to fine-tune promoter expression.

### caQTL-eQTL pairs are more often enriched for disease

Finally, to further explore the relationship between chromatin accessibility and gene expression levels, we then intersected genetic variants that were associated with both caQTL and eQTL effects in the same cell type (dual-QTLs). We found that for 93.0% of the eGenes at least one SNP drove both chromatin accessibility and expression levels (**Suppl. Fig. 6**). For the majority (78.0%) of the dual-QTLs, variants increasing chromatin accessibility levels were also associated with elevated gene expression levels, in line with previous observations derived from scMO data of 428 healthy Chinese individuals that showed 85.6% of the dual-QTLs being positive associations^29^ (**Fig. 4a**). We suspect the positive associations capture mostly activating regulatory processes (**Fig. 4b**), while the negative associations capture mostly repressive processes (**Fig. 4c**).

**Figure 4.**
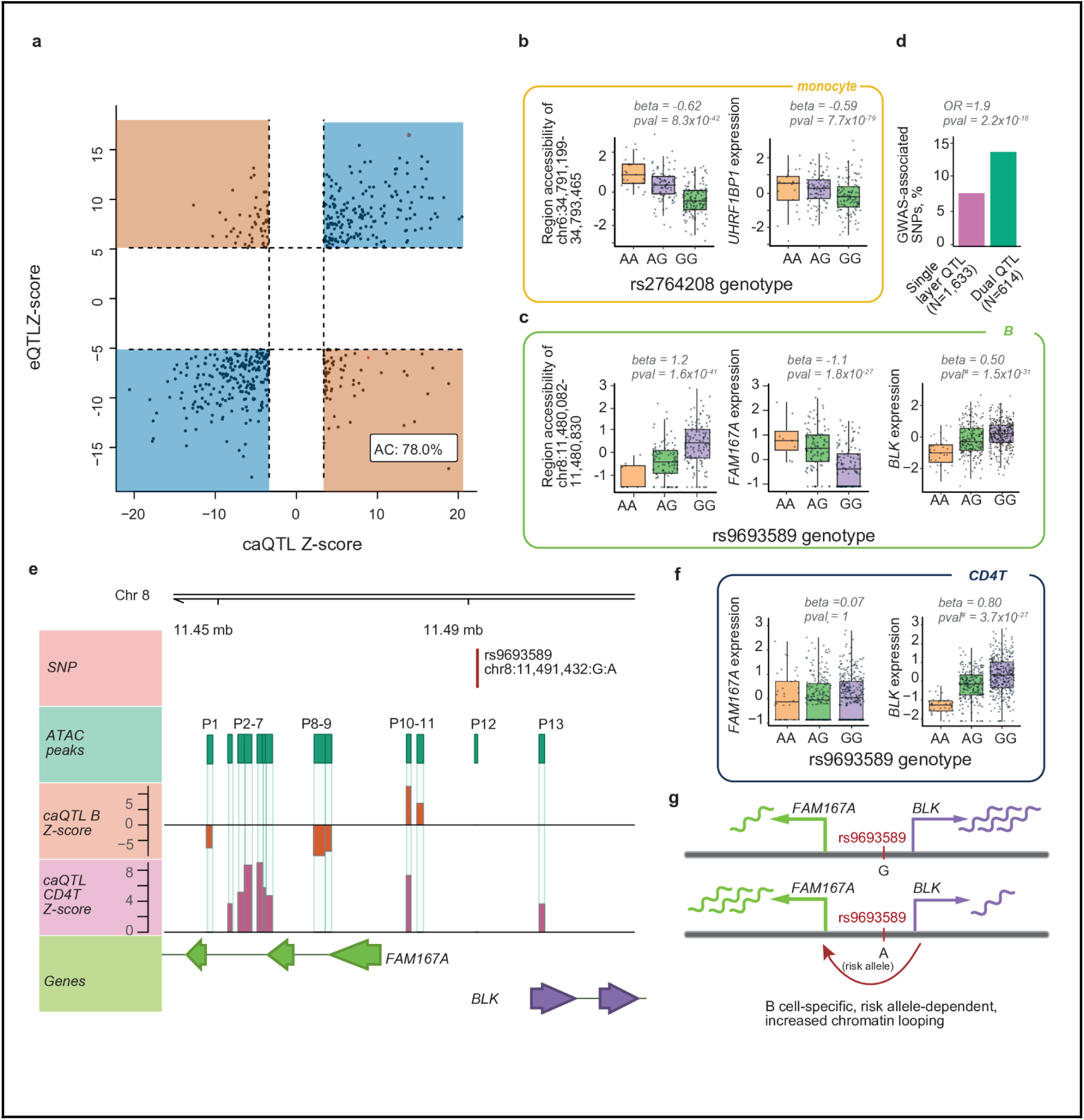
Dual-QTLs are enriched for disease. **a.** Scatterplot showing the eQTL and caQTL Z-score concordance. When multiple variants regulate the same peak-gene combination, we selected the variant with the strongest eQTL Z-score, after which the strongest matching caPeak was selected in the same cell type. **b.** Example of an immune disease-associated dual QTL (SLE) whose relationship between chromatin accessibility and gene expression is concordant **c.** or is associated with both a concordant (FAM167A eGene) and discordant effect (BLK eGene) (peak 10 in **e**). **d.** Immune disease GWAS enrichment of independent dual QTLs compared to single-layer acting QTLs (pruned by LD: R^2^>0.75). **e.** The FAM167A-BLK locus is a well known SLE-associated risk region in which genetic variants, among which the experimentally confirmed regulatory SNP rs9693589 (A risk allele), are associated with decreased BLK (in **d.** B and **f.** CD4+ T cells) and increased FAM167A (in **d.** B cells) expression. This reciprocal gene-gene regulation only occurs in B cells and coincides with several chromatin peaks only becoming more open in B cells (P1, P8-P9, P11). **g.** Proposed mechanism of action by which the allele-dependent, B cell-specific gene-gene regulation of the FAM167A-BLK locus can be explained. ^#^EigenMT multiple testing p-values are provided. EigenMT multiple testing correction was used to overcome permutation instability for the BLK gene.

Importantly, for these dual QTLs compared to single-layer acting QTLs we observed stronger enrichment for immune disease GWAS (OR = 1.9, Fisher’s Exact p = 2.2 x 10^-16^), in line with previous observations^30,31^ (**Fig. 4d**). As such, these dual QTLs can help us provide relevant insights in disease pathogenesis. For example, we found several dual QTLs that were associated with SLE risk, including a positive dual-QTL effect of rs2764208 in monocytes (**Fig. 4b**) and both negative and positive effects of rs9693589 in B cells (**Fig. 4c, 4e**). In the first example, the risk allele (G) at rs2764208 reduces both chromatin accessibility at peak chr6:34,791,199-34,793,465 and expression levels of *UHRF1BP1* (**Fig. 4b**). This is in line with previous observations in PBMCs of SLE patients that showed lower *UHRF1BP1* expression levels compared to healthy controls^31^. Mechanistically, *UHRF1BP1* is a binding partner of the epigenetic regulator *UHRF1*, whose loss has been shown to trigger type I interferon signalling through retrotransposon demethylation^32^ (Doyle et al., 2023). Together, this provides a regulatory explanation how a well-established SLE GWAS locus^33,34^ (Gateva et al. 2009; Zhang et al., 2011) can contribute to the increased interferon signaling that characterises SLE pathogenesis, with monocytes acting as key effector cells^35^ (Yao et al., 2023).

In the second example, the risk allele (A) at rs9693589 is associated with decreased chromatin accessibility at peak chr8:11,480,082-11,480,830 and increased *FAM167A* expression in B cells (**Fig. 4c**). This gene is thought to play a role in activation of the noncanonical NF-κB pathway^30^. Interestingly, this same allele also decreases *BLK* expression in both CD4+ T and B cells (**Fig. 4c, 4f**). These observations align with experimental evidence in B cells that shows that activating the risk variant region with CRISPR lowered BLK expression, and the variant showed allele-specific regulatory effects in a massively parallel reporter assay^36^. Importantly, the reciprocal gene regulation is only observed in B cells, which coincides with several chromatin peaks only becoming more open in the B cells (**Fig. 4e**) and risk allele-dependent increases in chromatin looping between *BLK* and *FAM167A* being previously reported in Hi-C data of LCL B cells (**Fig. 4g**)^36,37^. Together, this may explain how the downstream effects of this risk allele propagate differently in B cells, thereby contributing potentially to the enhanced survival of autoreactive B cells in SLE patients^35^.

Collectively, these examples illustrate the complexity of genetic regulation, and the importance of jointly considering chromatin accessibility and gene expression to understand the functional impact of non-coding disease-associated genetic variation.

### Genetic variants can rewire TF control of gene expression

While dual QTLs provide a way to link chromatin peaks to genes (**Fig. 4a**), this approach can only be applied when the same genetic variant regulates both chromatin and expression. Moreover, it remains difficult to distinguish causal (chromatin → expression) from consequential chromatin changes (expression → chromatin) without additional information. To overcome these limitations, SCENIC+^38^ uses single cell observations and integrates TF information to identify eRegulons (TF-CRE-target gene triplets) and distinguish causal from consequential chromatin changes. Applying SCENIC+ to our data, we identified 109,787 eRegulons, comprising 126 unique TFs, 41,768 peaks and 6,833 target genes (**Fig. 1e, Suppl. Table 6**). Within these eRegulons, we observed a median of 4 TFs per gene (**Fig. 5a**), and 6 peaks per gene (**Fig. 5b**). Comparing the TF-gene and CRE-gene pairs within these eRegulons with matched sets of control pairs, we observed greater-than-expected overlap with a curated database of protein-protein interactions (STRING database^39^: OR = 1.6, p = 2.75 x 10^-37^) (**Suppl. Fig. 7a**) and experimentally-derived (K562 HiC data^40^: OR = 2.5, p = 1.4 x 10^-126^) peak-gene interactions (**Suppl. Fig. 7b**), respectively. Moreover, we overlapped our CRE-gene pairs with those that were previously reported in PBMCs using REUNION, an alternative gene regulatory inference approach.^41^ This revealed that 96% of the overlapping CRE-gene pairs were concordant (**Suppl. Fig. 7c**) and that 96% of the TF-gene pairs were concordant (**Suppl. Fig. 7d**). Together, this supports the validity of the regulatory connections that we have identified.

**Figure 5.**
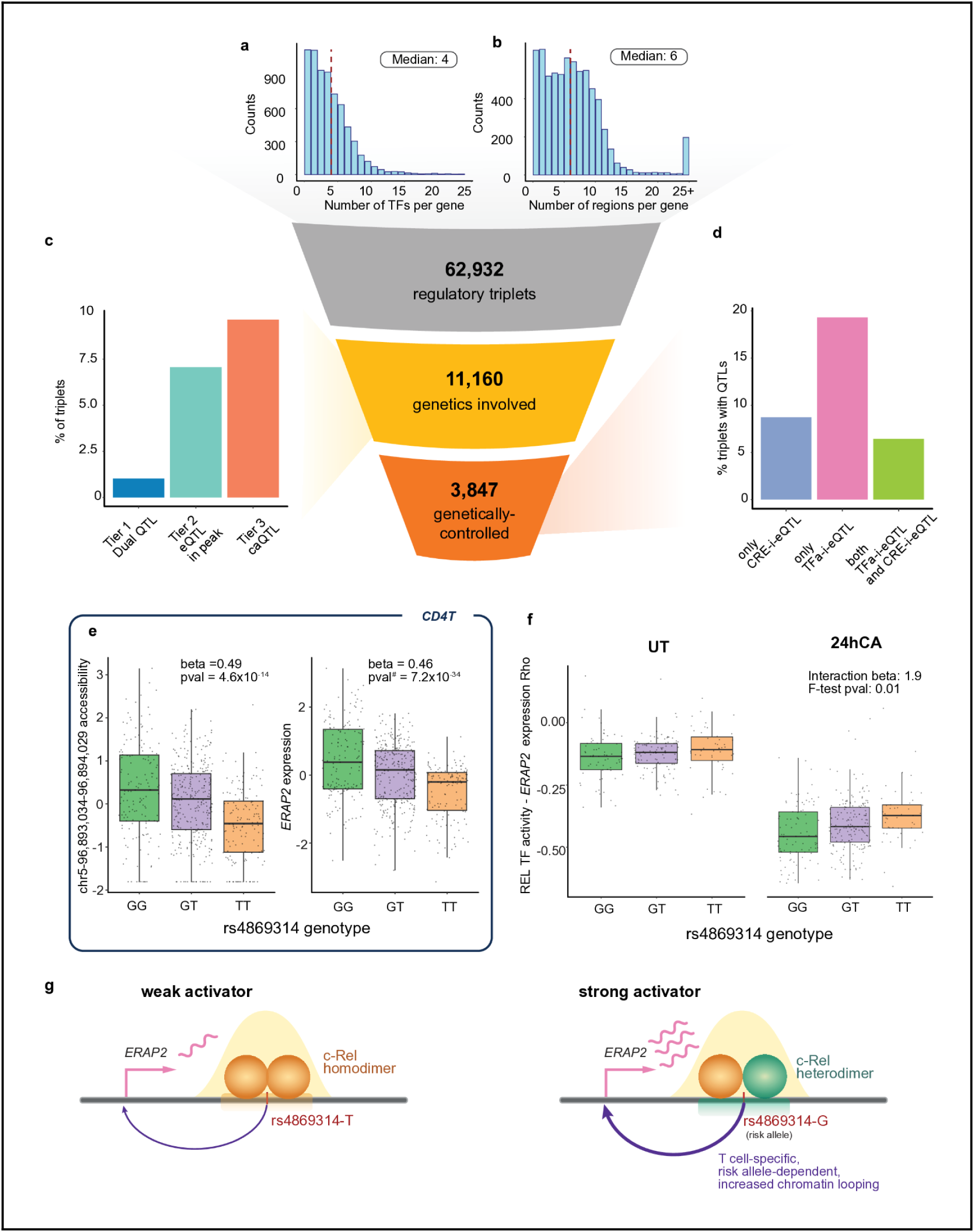
Genetic variants can rewire control of gene expression. **a.** Histogram depicting the number of TFs that are linked to the same target gene through SCENIC+-derived regulatory triplets. A median of 4 TFs are linked to the same target gene. **b.** Histogram showing the number of chromatin peaks that are linked to the same target gene through SCENIC+-derived regulatory triplets (eRegulon). A median of 6 peaks are linked to the same target gene. **c.** The proportion of SCENIC+ triplets as part of an eRegulon that overlap with genetic variants associated with a matching 1. dual-QTL (matching peak-gene), 2. eQTL within an accessible peak (matching peak-gene), or 3. caQTL (matching peak). Variants have not been classified in multiple tiers, and have been preferentially classified in a higher tier. **d.** The types of interaction-eQTLs with TF activity, chromatin accessibility or both. **e.** caQTL and eQTL regulated by SNP rs4869314 in CD4+ T cells for the region-gene pair that is part of the REL-ERAP2 eRegulon. **f.** Stimulation-specific interaction of REL-ERAP2 TF activity-i-eQTL. **g.** We expect the arthritis risk allele (G) of SNP rs4869314 to increase ERAP2 expression by promoting formation of a strongly activating c-Rel heterodimer complex at the enhancer under basal or CA-stimulated conditions. In contrast, the non-risk allele preferentially recruits a weakly activating c-Rel homodimer. As a result, changes in c-Rel TF activity produce a larger shift in regulatory complex composition in risk-allele carriers, i.e. from a strongly activating heterodimer toward a weakly activating homodimer, leading to a stronger negative relationship between c-Rel activity and ERAP2 expression. ^#^EigenMT multiple testing p-value is provided. EigenMT multiple testing correction was used to overcome permutation instability for the ERAP2 gene.

Next, we overlapped our eRegulons with the QTLs identified earlier (**Suppl. Table 2, 5**), and found that for 1.7% of the regulatory triplets genetics is involved (**Fig. 5c**). We grouped these genetic variants in 3 tiers reflecting their likelihood of influencing the regulatory triplet: 1.3% were overlapping a dual-QTL (tier 1) and 6.8% an eQTL (tier 2) that was matching the CRE-gene pair of an eRegulon. The remaining 8.9% overlapped with a caQTL whose caPeak was part of an eRegulon (tier 3). Potentially, the matching gene for these tier-3 CREs was not detected in our eQTL analysis because caQTLs are generally easier to identify than eQTLs: changes in chromatin accessibility can occur even when the downstream expression effect is too subtle to detect, or manifests only in cellular contexts not captured in our dataset^42,43^.

Next, we assessed whether these genetic variants can rewire the TF control of gene expression by testing these specific variant-regulatory triplet combinations in a linear regression model that explains target gene expression using TF activity and the specific genetic variant (TFa-i-eQTL). For the TF activity we used the AUCell target gene score per cell from SCENIC+ (**Methods**). Importantly, for the TFa-i-eQTLs that were initially significant, we assessed possible circularity by repeating the same analysis but now excluding the eGene as target gene from the TF activity calculation. This step invalidated 2% of our initially discovered TFa-i-eQTLs, leaving 3,857 effects (B&H FDR < 0.05) (**Suppl. Table 7**). These remaining effects represent the final TFa-i-eQTL set, which represent 25.7% of all the regulatory triplets where genetics is involved (**Fig. 5d**). If instead of TF activity the same linear model was run, but now using CRE peak accessibility or TF expression (TFe), 15% and <1% of the genetically-controlled regulatory triplets were annotated as CRE-i-eQTL or TFe-i-eQTL (B&H FDR < 0.05), respectively (**Suppl. Table 8**)—with the far majority (96.1% of the CRE-i-eQTLs) also identified as TFa-i-eQTL with the same effect direction (**Fig. 5d**, **Suppl. Fig. 8a, 8b**). While theoretically we expect that all TFa-i-eQTLs would also be a CRE-i-eQTL, the larger sparsity of the CRE accessibility compared to the TF activity measurement likely reduced the power to detect CRE-i-eQTLs. In the case where the limiting factor is the chromatin openness and not the TF abundance, it could still be possible a CRE-i-eQTL is detected, but a TFa-i-eQTL is not. Similarly, we noticed that TF expression and TF activity generally correlate well (**Suppl. Fig. 9a**), but there are exceptions. Especially upon stimulation, we observed a generally better correlation between TF expression and activity (Wilcoxon signed rank p < 1.3 x 10^-14^) (**Suppl Fig. 9b**). This may be especially true for stimulation-responsive TFs that are already expressed at baseline, allowing their activity to be triggered immediately upon stimulation through rapid nuclear translocation^44–48^. Together, this showcases how using TF activity instead of CRE accessibility or TF expression can better capture genetically controlled gene regulatory relationships while also overcoming the reduced statistical power due to sparsity of the data.

Finally, to showcase how combining our QTLs with SCENIC+ regulatory triplets can help to improve our disease understanding, we highlight SNP rs4869314 whose risk allele (G) is associated with juvenile idiopathic arthris^49^. The risk allele was associated with increased *ERAP2* expression (in all major immune cells) and increased accessibility of the peak in which the SNP resides (chr5:96,893,034-96,894,029) (specifically in CD4+ T cells) (**Fig. 5e, Suppl. Table 2, 5**). Previously, H3K27Ac HiChIP analysis indicated that the region harboring this variant physically interacts with the *ERAP2* promoter, and that the risk allele increases the strength of this interaction in T cells^50^. This suggests that by altering this promoter-enhancer interaction, the variant may influence which regulatory complexes assemble at the *ERAP2* promoter.

Overlap with SCENIC+ eRegulons, indicated that the TF c-Rel may be the regulator binding in this enhancer region, thereby regulating *ERAP2* (**Suppl. Table 6**). c-Rel is part of the NF-κB TF family. These TF family members all have closely related binding motifs, and c-Rel can form homo- or heterodimers with other members of the family (p50, p52, RelA, RelB^51,52^c-rel). Depending on its interacting partners, c-Rel normally acts as either a strong (c-Rel:p50) or weak activator (c-Rel:p52, c-Rel:c-Rel). Explicit modelling of the relationship between c-Rel TF activity, the *ERAP2* risk SNP, and *ERAP2* expression using TFa-i-eQTL mapping (**Suppl. Table 7**) indicated that the risk allele increases the repressive capacity of c-Rel on *ERAP2* expression (**Fig. 5f**). At first sight, this appears contradictory to the observation that the risk allele is associated with higher *ERAP2* expression at baseline (**Fig. 5e**). However, a genotype-dependent shift in TF binding at this locus may reconcile these findings.

Specifically, we propose that the SNP alters the relative binding preference between c-Rel and another NF*-κ*B family member (**Fig. 5g**). For the non-risk allele, c-Rel likely binds efficiently and drives moderate *ERAP2* activation. In contrast, the risk allele may reduce c-Rel binding affinity, thereby allowing stronger NF*-κ*B activators to occupy the site more frequently. As a result, individuals carrying the risk allele exhibit higher *ERAP2* expression under basal conditions. Yet, when c-Rel activity increases, the displacement of these stronger activators by c-Rel leads to a disproportionately larger reduction in *ERAP2* expression, consistent with the observed stronger negative c-Rel effect in risk-allele carriers (**Fig. 5f**). This proposed mechanism by which c-Rel regulates *ERAP2* is previously seen for TNF-α-induced RelA-dependent inflammatory genes^53^. Therefore, it is not unlikely that related regulatory mechanics could be involved in the regulation of *ERAP2*.

Normally, *ERAP2* is involved in the trimming of intracellular peptides prior to their loading onto HLA class I molecules^54^. Therefore, excessive *ERAP2* activity can lead to over-processing of self-derived peptides, disrupting normal antigen presentation^55–57^ and potentially generating epitopes that activate autoreactive CD8*⁺* T cells. Together, this could contribute to the chronic inflammation observed in the joints of arthritis patients.

Summarized, by leveraging scMO measurements we could obtain less sparse and likely more accurate estimates of single-cell TF activity than would be possible using TF expression or chromatin accessibility alone. This enabled us to study how genetic variants can rewire TF control of gene expression, ultimately shaping inter-individual variation in disease risk.

## Discussion

Gene expression alone can only explain up to 40-50% of all GWAS loci^6,7^. It has been suggested that context-specificity^58,59^ or intrinsic differences between GWAS and eQTL loci may underlie this gap^60^. Here we explored whether looking at the broad picture of gene regulation could provide a more complete understanding of these GWAS loci. For this we applied same-cell single-cell multi-ome analyses within 264 donors in both pathogen-stimulated and unstimulated conditions. We suspected that considering both epigenetic and expression data at the same time could improve identification of the causal variants, as you would expect chromatin to be open in the context in which the eQTL effect takes place. For 42% of our lead eQTL variants chromatin was open in the matching condition. Moreover, variants that controlled both chromatin accessibility and gene expression were almost twice as often associated with disease than those acting on only a single data layer, supporting our hypothesis. This aligns with previous findings from the iPSC Omics Resource, in which variants that act on chromatin accessibility, gene expression levels and H3K27ac showed the largest disease enrichment^61^.

Then focussing on stimulation-dependent effects, we noticed these were more often located further away from the gene, mostly in distal enhancers. This overlaps with the notion that context-dependent gene regulation is often controlled by enhancers, whereas more general gene activity is controlled by promoters^62,63^. Moreover, this more distal location of stimulation-dependent effects is a pattern that is also seen for GWAS loci^60^, indicating that such context-dependent effects can provide additional explanation of GWAS loci^58–60^.

To explore the regulatory mechanisms driving our (stimulation-dependent) eQTLs, we employed the enhancer-driven GRN inference tool^38^. This led to the identification of 109,787 regulatory triplets that were identified by correlating TF activity to target gene expression, and to candidate enhancer regions. These TFs are identified through motif enrichment analysis in the co-accessible candidate enhancer regions. As such, one of the shortcomings of this approach is that it is inherently bounded by the completeness of TF motif databases: fewer than 50% of the over 1,600 human TF-encoding genes have a validated binding motif in resources such as JASPAR^64^ or HOCOMOCO^65^, and transcriptional co-regulators acting without direct DNA contact remain entirely invisible to motif-scanning pipelines. Emerging motif-agnostic tools directly address this limitation:REUNION^41^ infers peak-TF-gene triplets without relying on motif detection, recovering binding associations in regions lacking a canonical motif and improving target gene expression prediction. Epiregulon^66^ similarly handles co-regulators without defined motifs through a co-occurrence-based framework of TF expression and the accessibility of the TF’s ChIP-supported binding sites, while *de novo* approaches such as DENIS^67^ identify uncharacterised footprints in ATAC-seq data in a cell-type-specific manner.

Another shortcoming of the SCENIC+ approach, is that genetic relationships cannot be modelled explicitly. Previous studies have shown that not just gene expression, but also its regulation can be genotype-dependent^10,47,68,69^, underscoring the importance of considering genetic variants to improve the identification of regulatory triplets. By overlapping our caQTLs and eQTLs with the SCENIC+ regulatory triplets, and then explicitly modeling genotype-dependent TF activity- or CRE-to-eGene interactions, we showed that about 4% of our regulatory triplets were genetically controlled. This indicates that while SCENIC+ does not explicitly consider genetic variants, it can still identify triplets that are controlled by genetic variation. We expect that triplets that are now missed will be enriched for target genes that are controlled by variants with a high MAF or where the variant has a large effect size on the target gene. This is because such variants influence the TFactivity-to-target-gene relationship in a substantial fraction of cells, making the true regulatory relationship noisier. To not miss triplets that are under genetic control, future GRN inference tools should explicitly model this genetic variation, such as is implemented in Epiregulon, which explicitly models how shifts in TF activity can impact the genes it regulates. A shift in target gene regulation can occur due to 1) genetic mutations in the TF itself, 2) changes in the TF’s interaction partners, or 3) changes in the cellular context causing the TF to bind to a different set of genomic regions. Focussing on the impact of genetic variation in our results, we have seen putative cases of both the second (e.g. shift between weak and strong c-Rel regulatory complex formation, **Fig. 5g**) and third scenario (e.g. genetic variant impacting the binding affinity of the c-Rel TF, **Fig. 5g**). This indicates that by also incorporating genetic information will allow us to identify many more regulatory triplets and gain a more complete view of gene regulation.

Beyond these limitations, a key contribution of this study is the prioritization of genetic variants that influence gene regulation. While we identify variants that overlap or interact with regulatory elements and associate with changes in TF activity and gene expression, these statistical associations alone cannot confirm that a given variant causally drives the regulatory effect it is linked to. The resulting variant lists should be viewed as a starting point for hypothesis generation and as a foundation for targeted experimental follow-up, rather than a definitive causal map. This experimental follow-up could consist of approaches such as massively parallel reporter assays to screen regulatory activity, CRISPRi perturbations to establish causal gene links, and motif analysis to identify disrupted TF binding events. Such a strategy has recently successfully been applied in primary B cells for fine-mapped eQTL variants that colocalize with GWAS risk loci^36^. Applying these strategies more broadly also to the variants prioritized here, represents a natural next step toward establishing the mechanistic basis of immune gene regulation and disease susceptibility.

Together, our work demonstrates that combining sc-multiome data across conditions with enhancer-driven GRN inference and variant prioritization provides a deeper, more mechanistic understanding of immune gene regulation. This integrated framework can help explain more GWAS loci and nominates context-specific regulatory variants for follow-up experiments. As population-scale single-cell cohorts grow and motif-agnostic, genotype-aware inference tools mature, we anticipate that this integrative framework will increasingly resolve the full genetic and context-dependent architecture underlying complex immune traits and disease susceptibility.

## Methods

### Ethics approval and informed consent

The LifeLines DEEP^70^ study was approved by the ethics committee of the University Medical Centre Groningen, document number METC UMCG LLDEEP: M12.113965. All participants signed informed consent from prior to study enrollment. All procedures performed in studies involving human participants were in accordance with the ethical standards of the institutional and/or national research committee and with the 1964 Helsinki declaration and its later amendments or comparable ethical standards.

### PBMC collection

Whole blood from X European white background individuals of the northern Netherlands population cohort Lifelines Deep^70^, Lifelines Next^71^ and LongCOVID was drawn into EDTA-vacutainers (BD). PBMCs were isolated and maintained, as described previously^10^. In short, PBMCs were isolated using Cell Preparation Tubes with sodium heparin (BD) and were cryopreserved until use in RPMI1640 containing 40% FCS and 10% DMSO. After thawing and a 1 h resting period, cells were washed twice in a medium supplemented with 0.04% bovine serum albumin and directly processed for scRNA-seq. Cells were then counted using a hemocytometer, and cell viability was assessed by Trypan Blue.

### Library preparation

Libraries were prepared with 10x Chromium Next GEM Single Cell Multiome ATAC + Gene Expression workflow (CG000365) V2 chip and V2 reagents. Eight samples were pooled per 10x lane, in duplo, aiming for a total recovery of 10,000 cells per lane. Four lanes per chip were loaded, aiming at an expected yield of 2500 cells per sample.

### Genomic DNA isolation and imputation

Genomic DNA was isolated from unused nuclei remaining after preparation of the nuclei suspension from the 10x workflow. Genomic DNA was purified using the Wizard SV Genomic DNA Purification System (Promega) according to the manufacturer’s instructions. Genotype data was obtained with Global Screening Array, as performed at Rotterdam MC. This data was subsequently imputed using the 1000G reference with the sc-eQTLgen pipeline v1.0^25^.

### Single cell sequencing and data preparation

Sequencing was done at BGI (Hong Kong) on an MGI2000 sequencer, featuring 100bp paired-end reads. Demultiplexing was performed at BGI. Cellranger ARC v2.0 was used to perform filtering, deduplication, barcode counting, and alignment of both gene expression and open chromatin data, using the human GRCh38-3.0.0 genome as a reference.

### Demultiplexing and sample assignment

Souporcell^17^ v2 was used based on the gene expression data, to define genotypic clusters for each of the 10x sample pools, using only locations on the genome defined by Souporcell as showing common variation. Cells defined as doublets instead of clusters were removed. Genotypes constructed for each cluster in Souporcell, were correlated to the Global Screening Array genotypes present in that 10x sample pool to assign the genotypic clusters to individuals.

Freemuxlet^72^ was used based on the open chromatin data for each of the 10x sample pools.

### Ambient RNA correction

Cellbender^16^ v3 was used on the raw gene count matrices for each sample pool coming from cellranger ARC, using the default method of estimating the correct number of true cells dynamically, to remove gene counts coming from ambient RNA that were incorrectly assigned to cells.

### Single Nucleus Gene expression QC

Ambient-RNA corrected count matrices were merged across sample pools. Median Absolute Deviation was then calculated for the number of UMIs and number of unique genes of each cell. Cells with less than 200 total UMIs or more UMIs more than 3 MADs were removed, as well as cells with less than 200 different genes expressed or more unique genes than 3 MADs.

### Chromatin Accessibility quantification

Fragment counts from the cellranger ARC output were modelled to represent fragment counts instead of read counts^73^.

These fragments were subsequently aligned to a reference bed file^24^ containing areas of the human that can exhibit open chromatin, and the number of fragments aligning to these areas were counted per nucleus.

### Single Nucleus Chromatin Accessibility in Signac

Fragment count matrices per sample pool calculated from the re-aligment to the reference bed file were loaded into Signac and combined across pools. Cells with less than 3000 or more than 30000 counts were removed. Based on the cellranger ARC output, cells were removed where less than 50% of their reads fell into peaks. Next, cells were removed with a nucleosome signal larger than 4 or TSS enrichment score of less than 3. Finally, sample assignment and doublet information was added stemming from the expression data, and cells annotated as doublets were removed.

The data was normalized using term frequency inverse document frequency, after which partial singular value decomposition was run on the transformed counts. LSI dimensional reduction was performed, followed by KNN-clustering and further dimensional reduction using UMAP. Cell type annotations were added based on the expression data, assigning cells not present in the expression data to the cell type most prevalent in the cluster the cell was assigned to. The data was then split up per cell type for further data analysis.

### Cell type assignment

A PBMC 10x multi-ome of 10K cells dataset was downloaded using the SeuratData package. This dataset was subsequently filtered to only contain cells that had both RNA and ATAC measurements. The RNA data was normalized using the total number of UMIs per cell, and subsequently transformed using natural log. Principal components were calculated over the transformed counts. The ATAC data was normalized using term frequency inverse document frequency, after which partial singular value decomposition was run on the transformed counts. A weighted nearest neighbours graph was subsequently calculated with the ‘FindMultiModalNeighbors’ function using the first 30 principal components of the RNA data, and the second to third LSI components of the ATAC data. This neighbour graph was used to construct a 2D Uniform Manifold Approximation and Projection of the data, and as input for clustering using the smart local moving algorithm with a resolution of 2.0.

Data was processed for each of the 80 sample pools separately, using Seurat to create an object with cellbender corrected RNA counts, and ATAC counts based on the ATAC fragments from cellranger ARC. Each sample pool was then processed as described for the reference dataset listed before. Seurat’s Azimuth method was finally used to project each sample pool onto the multimodal reference dataset listed before, transferring cell type annotations to the query sample pool. Cell type annotations were then aggregated and transferred to the object containing all sample pools.

The expression data across all 80 sample pools was then normalized using the SCTransform method in Seurat, after which Principal Component Analysis was performed and the cells were clustered based on the first 30 principal components. To annotate cells that did not have chromatin data with a cell type, each cell in a cluster was assigned the cell type most prevalent in that cluster.

### Differential Gene Expression Analysis

Expression was pseudobulked by aggregating the counts for each unique combination of donor, pool, cell type and stimulation condition. Pseudobulks derived from less than five nuclei or having less than 10000 total UMIs were discarded. Pseudobulks were then normalized using TMM normalization. Limma dream^74^ was used to model the stimulation status as dependent variable, while correcting for age, sex as fixed effects and sample pool and donor as random effects, weighting each pseudobulk by the number of nuclei it was generated from. Raw P-values for the stimulation condition were Bonferroni corrected using the number of genes. A bonferroni-corrected P-value of 0.05 was considered significant.

### Differentially Accessible Regions

Pseudobulk ATAC measurements were constructed by aggregating across each combination of the stimulation condition, donor, pool and cell type. Pseudobulks derived from less than five nuclei or having less than 10000 total UMIs were discarded. DARs were only tested if the average absolute LFC was >0.1. Pseudobulks were then normalized using TMM normalization. Limma dream^74^ was used to model the stimulation condition as dependent variable, while correcting for age, sex as fixed effects and sample pool and donor as random effects, weighting each pseudobulk by the number of nuclei it was generated from. Raw P-values for the stimulation condition were Bonferroni corrected using the number of regions. A bonferroni-corrected P-value of 0.05 was considered significant.

### Topic modelling

Fragment matrices were exported from Signac and loaded separately for each cell type and subsequently loaded into pycistopic^75^. A binary matrix was then constructed from the fragment matrix, replacing all values above zero with one. Regions present in less than 0.1% were removed from the analysis. Topics were then modelled using the Otsu method, with the number of topics set to 120. Cells were then binarized to topics and accessibility was imputed using the topic information. Cell to topic assignments and imputed accessibility was finally exported.

### eRegulon detection

Signac objects after quality control were converted to Scanpy objects per cell type and used in conjunction with the pyCistopic objects per cell type as input for the SCENIC+ pipeline^38^. A precomputed database of TF-gene based on was used, as well as the database linking genomic patterns to transcription factors.

### eQTL mapping

The single-nucleus expression data was first normalized by calculating the total UMI per nucleus, then dividing total UMIs per nucleus by this mean to get a scaling factor, then applying this scaling factor to all gene counts for each nucleus^76^. Expression was next pseudobulked by summing the expression for each combination of cell type and sample. The first 20 principal components were then calculated over these pseudobulks for each cell type separately. We then fit a linear mixed model using the sc-eQTLgen QTL mapping tool LIMIX-QTL^25^, predicting gene expression by genetic variants, while correcting for the first 20 expression PCs and stimulation status as fixed effects, and the donor assignments of the samples as random effect. We confined our analysis to cis-eQTL effects with a windows size of 1M. We used 1000 variant-level permutations for each gene to control for false discovery at the variant level, and then performed q-value correction^77^ over the strongest permutation-level effects per gene to account for gene-level false discovery. We considered genes with a gene-level qvalue of <0.05 significant associations.

### caQTL mapping

The single-nucleus chromatin accessibility data was normalized and pseudobulked in the same way as the gene expression data as described in the We confined our analysis to cis-caQTL effects with a windows size of 50k. We used 20 variant-level permutations for each chromatin region to control for false discovery at the variant level, and then performed q-value correction^77^ over the strongest permutation-level effects per chromatin region to account for chromatin-region-level false discovery. We considered chromatin regions with a chromatin-region-level qvalue of <0.05 significant associations.

### Processing of LCL dataset

To replicate our single-cell chromatin accessibility QTLs, we leveraged LCL B cell data from 100 British samples from the 1000 Genomes Project. Specifically, we used the processed and normalized bulk ATAC-seq log-transformed FPKM values from Kumasaka et al^27^.

Genotype data were obtained from the high-coverage whole-genome sequencing (WGS) data of the 1000 Genomes Project (GRCh38), generated by Byrska-Bishop et al^78^. The overlap between donors with WGS data and those with ATAC-seq measurements comprised 91 individuals.

For replication of the caQTL signal, we lifted over the peaks by Kumasaka et al to hg38 by using the UCSC liftover tool^79^. We then intersected our peaks with the LCL peaks, considering peaks that overlapped with at least 1 bp as the same genomic region. We then mapped caQTLs using a linear model implemented in LIMIX QTL^25^, including SNPs with a MAF > 1%, and HWE P > 0.0001.

Leveraging this setup, we considered our identified single cell caQTLs to be replicated, when reaching nominal significance and a consistent effect direction.

### Processing of OneK1K dataset

To replicate our single-nucleus expression QTLs, we leveraged summary statistics obtained before meta-analysis in sc-eQTLgen^80^. We then performed q-value correction^77^ over the strongest permutation-level effects per gene to account for gene-level false discovery. We considered genes with a gene-level qvalue of <0.05 significant associations.

### stimulation-interaction-eQTL mapping

Single-nucleus expression data was processed as described in ‘eQTL mapping’. We selected genes showing a significant eQTL result for interaction-eQTL mapping in the corresponding cell type. We then fitted a linear mixed model using LIMIX-QTL^25^, predicting gene expression by genetic variants, while correcting for the first 20 expression PCs and stimulation status as fixed effects, and the donor assignments of the samples as random effect. We then used genotype*stimulation status as an interaction term. To control for false discovery in the interaction-eQTL mapping, we made use of eigenMT. In short, eigenMT performs eigenvalue decomposition of the genetic variants tested for a feature, then performs bonferroni based on the nominal p values for that feature, based on the number of eigenvalues required to explain 98% of the variance. We subsequently selected the top effect per feature, and performed q-value correction^77^ to control for false discovery at the gene level. We considered genes with a qvalue<0.05 to be significant stimulation-dependent interaction-eGenes.

To investigate if the distance between variants and eQTL genes was significantly different for stimulation-interaction-eQTLs vs non-interaction-eQTLs, we matched top interaction-eQTLs to non interaction-eQTLs with an baseline eQTL effect size, gene body size and MAF within 5% of the interaction-eQTLs. We then fit a linear regression model explaining the distance by the presence of an interaction effect, correcting for the same categories we matched on.

### stimulation-interaction-caQTL mapping

Single-nucleus chromatin data was processed in the same way as expression data, as described in ‘stimulation-interaction-eQTL mapping’.

### statistical fine-mapping

QTL summary statistics coming from the interaction-QTL mapping were filtered on features which were qvalue<0.05 significant in the condition-independent QTL mapping. The 1000G^81^ high-coverage European population was used as an LD reference panel. Finemapping was then performed per cell type, per feature using SuSIE^82^ with an initial maximum of causal signals per locus set at 10, and 100 interactions. When a feature did not converge, the finemapping was retried with double the iterations, a max number of three times. If after three retries convergence is still not achieved, the maximum number of causal signals is halved until it reaches one, where the number of detected causal signals is set to be one.

### TF activity-interaction-eQTL mapping

Regulatory triplets identified in SCENIC were overlapped with output from eQTL mapping by first filtering the mapping results to only contain variant-gene associations where the genetic variant was present in the chromatin region linked to the gene through SCENIC+. The top effect per gene per cell type was then selected to be tested. Additionally, caQTL effects were filtered for the chromatin regions identified associated with genes in SCENIC+. The top effect per region per cell type was then selected among the filtered caQTLs. The eRegulon (describing TF activity) for these regions and genes was then finally selected from the SCENIC+ results.

TF activity per cell as gene AUCs for these eRegulons was extracted from the SCENIC+ h5mu object, exported as a cell by tf activity matrix and each eRegulon was transformed to fit a gaussian distribution. Cell by gene expression data of cells present in the cell by activity matrix was also transformed to fit a gaussian distribution. A linear mixed model was then fitted where expression of the gene is explained by the TF activity, while correcting for total expression of a cell and eQTL genotype as fixed effects, with experimental batch and donor as random effects. Next an interaction model was fit with genotype * TF activity as an interaction term. Residuals of the first and interaction model were compared using a two-sided F-test. Interaction- and F-test p-values were finally corrected using Benjami-Hochberg correction to control for false discovery. Interactions where both the corrected interaction- and F-test p-values were smaller than 0.05 were considered significant.

To control for possible confounding effects where the target gene was part of the gene set from which the TF activity was calculated, for the significant TFa-i-eQTLs, the TF activity was recalculated using the same AUCell method, with the TFa-i-egene removed from the gene set. The interaction analysis was then repeated using these recalculated TF activities. Associations surviving this second pass were considered true effects.

### CRE-interaction-eQTL mapping

Regulatory triplets were overlapped with eQTLs and caQTLs as described in ‘TF-interaction-eQTL mapping’. Expression data is made to fit a gaussian distribution, while accessibility data is binarized as [accessibility >1] = 1. A linear mixed model was fitted where the expression is explained by the openness of the chromatin region associated to the gene, while also correcting for total expression, and eQTL genotype as fixed effects, and experimental batch and donor as random effects. Next an interaction model was fit with genotype*accessibility as an interaction term. Residuals of the first and interaction model were compared using a two-sided F-test. Interaction- and F-test p-values were finally corrected using Benjami-Hochberg correction to control for false discovery. Interactions where both the corrected interaction- and F-test p-values were smaller than 0.05 were considered significant.

## Ethics declaration

### Competing Interest Statement

L.F. has ongoing contract-based research with Biogen and Roche, not related to this work.

### Funder Information Declared

ZonMw, The Dutch Organisation for knowledge and innovation in health, healthcare and well-being, https://ror.org/01yaj9a77, 09150182010019, 10430302110002, 09150172310068 Dutch Research Council, 223.041 Oncode Institute

### Author contributions

Conceptualisation: M.Wi., L.F., R.O., M.K.

Data generation: J.N., M.Wi.

Data curation & Analysis: R.O., M.K., D.K., M.We

Funding acquisition: M.Wi., L.F.

Methodology and software: R.O., M.K., M.J.B.

Supervision: M.W., L.F.

Writing – original draft and Visualisation: M.Wi., M.K., R.O.

Writing – review & editing: L.F., M.J.B.

## Supporting information

ST1 - cell QC

ST2 - eQTLs

ST3 - DEGs

ST4 - DARs

ST5 - caQTLs

ST6 - triplets

ST7 - TF-i-eQTLs

ST8 - CRE-i-eQTLs

## Acknowledgements

We are very grateful to all the volunteers who participated in this study. This work was carried out on the computer cluster of the Genomics Coordination Center, hosted at the University of Groningen Center for Information Technology.

L.F., D.K., M.J.B. are supported by a grant from the Dutch Research Council (ZonMW-VICI 09150182010019 to L.F. ZonMW LongCOVID grant 10430302110002), European Union’s Horizon Europe Research and Innovation Program grant 101057553 (LongCovid) and through a Senior Investigator Grant from the Oncode Institute and a grant from Saxum Volutum (Pericode). M.J.B. is supported by a grant from the Dutch Research Council (ZonMW-VIDI 09150172310068). M.W. is supported by a grant from the Dutch Research Council (Vidi 223.041)

## Data availability

Full summary statistics for Differential Gene Expression, Differential Accessibility, eQTLs, caQTLs, stim-interaction-eQTLs, stim-interaction-caQTLs are available via the companion web page at eqtlgen.org/sc/datasets/scmo.html. Processed expression and accessibility are also available here. Raw expression, raw accessibility and genotype data will be made available via LifeLines data management, also linked on the webpage.

## Code availability

The original code for the sc-eQTLgen imputation pipeline v1 (https://github.com/sc-eQTLgen-consortium/WG1-pipeline-QC), Souporcell v2 (https://github.com/wheaton5/souporcell), Seurat v5 (https://github.com/satijalab/seurat), LIMIX-QTL (https://github.com/single-cell-genetics/limix_qtl), SuSIE (https://github.com/stephenslab/susieR) and SCENIC+ (https://github.com/aertslab/scenicplus) are available via GitHub. All custom-written code is also available via GitHub (https://github.com/molgenis/wijst_multiome).

## Notes

https://eqtlgen.org/sc/datasets/scmo.html

### Summary of Updates

Updated figure one to show the filtered triplets number instead of the unfiltered triplet number

