## Supplementary Information for "Cell-specific regulatory circuits connect genetic variation to disease susceptibility"

### Suppl. Note 1

**Filtering of called consensus peaks**
We quantified the chromatin accessibility in each cell by overlapping our ATAC reads with peaks derived from a human reference set of consensus open chromatin regions (cPeaks)[1]. As a consequence, overlap of a single ATAC read with this reference would result in a peak call. Therefore, to prevent that background ATAC reads would negatively impact our peak calling, we restricted our further analyses to regions with sufficient coverage in that cell type or condition. To define this cut-off, we compared (before and after filtering): the 1. total number of peaks retained, 2. percentage of housekeeping peaks retained, 3. cell type-specificity of the peaks, 4. relative fraction of promoter and enhancer peaks, 5. replication with IHEC bulk monocyte called peaks[2]. We observed that applying filtering thresholds keeping only peaks that were present in at least 0.1% or 1% of the cells of a particular major cell type, retained 61.9% (777,433 unique peaks) and 17.7% (222,497 unique peaks) of the non-filtered peaks, respectively (**Suppl. Note Fig. 1a**). Importantly, despite filtering out many peaks, we barely lost any housekeeping peaks using the 0.1% (32 housekeeping peaks lost, 0.1% total housekeeping peaks) or 1% (151 housekeeping peaks, 0.6% total housekeeping peaks) filter. This indicates that these filtering thresholds barely affect those peaks that should be present in every cell type (i.e. the housekeeping peaks).

As we preferred to keep as many peaks as possible, while capturing relevant biology, we then assessed whether the 0.1% filter can strike this balance. Interestingly, while most peaks were shared across lineages before filtering, lineage-specificity became much more apparent after filtering (**Suppl. Note Fig. 1b**). On top of that, we noticed that the cPeaks that overlapped promoter and enhancer annotated ENCODE SCREEN peaks (PLS, pELS, dELS)[3], which are most relevant for creating gene regulatory triplets, are relatively enriched after applying the 0.1% filter (**Suppl Note Fig. 1c**).

Finally, we used an independent bulk monocyte ATAC-seq dataset of 33 samples from the IHEC resource[2] to assess how well our cPeaks overlap with this dataset, and define whether we can capture some of the expected cell type-specificity using this data. For this, we examined the overlap between our monocyte- and CD4+ T cell-called cPeaks and those IHEC bulk monocyte ATAC peaks that were present in at least 2 samples (to ensure comparing biological relevant signals that are also expected to be found in our samples). Before filtering, we did not see any difference in overlap with IHEC monocyte peaks for either our CD4+ T- or monocyte-called cPeaks. However, after applying the 0.1% filter to our cPeak-called peaks, we retained a similar level of overlap for the monocyte-called cPeaks (96.9%, OR = 42.2, Fisher's exact p-value = 0), whereas we lost substantial overlap of monocyte-specific peaks in a similar size cell type from the other lineage (CD4+ T cells, 82.3%, OR = 1.2, Fisher's exact p-value = $1.85 \times 10^{-227}$). The two associations differed significantly (log OR difference = 3.54, Z = 75.68, p-value = 0; **Suppl. Note Fig. 1d**).

Altogether, based on these analyses we kept peaks that were detected in at least 0.1% of the cells of a particular major cell type, as that gave the best balance between peak sensitivity and biological (i.e. driven by lineage-specificity, remain housekeeping peaks) over technical signal (i.e. driven by statistical power). Moreover, applying such a filter also discarded those peaks for which we would not have sufficient numbers of observations for any of our downstream analyses.

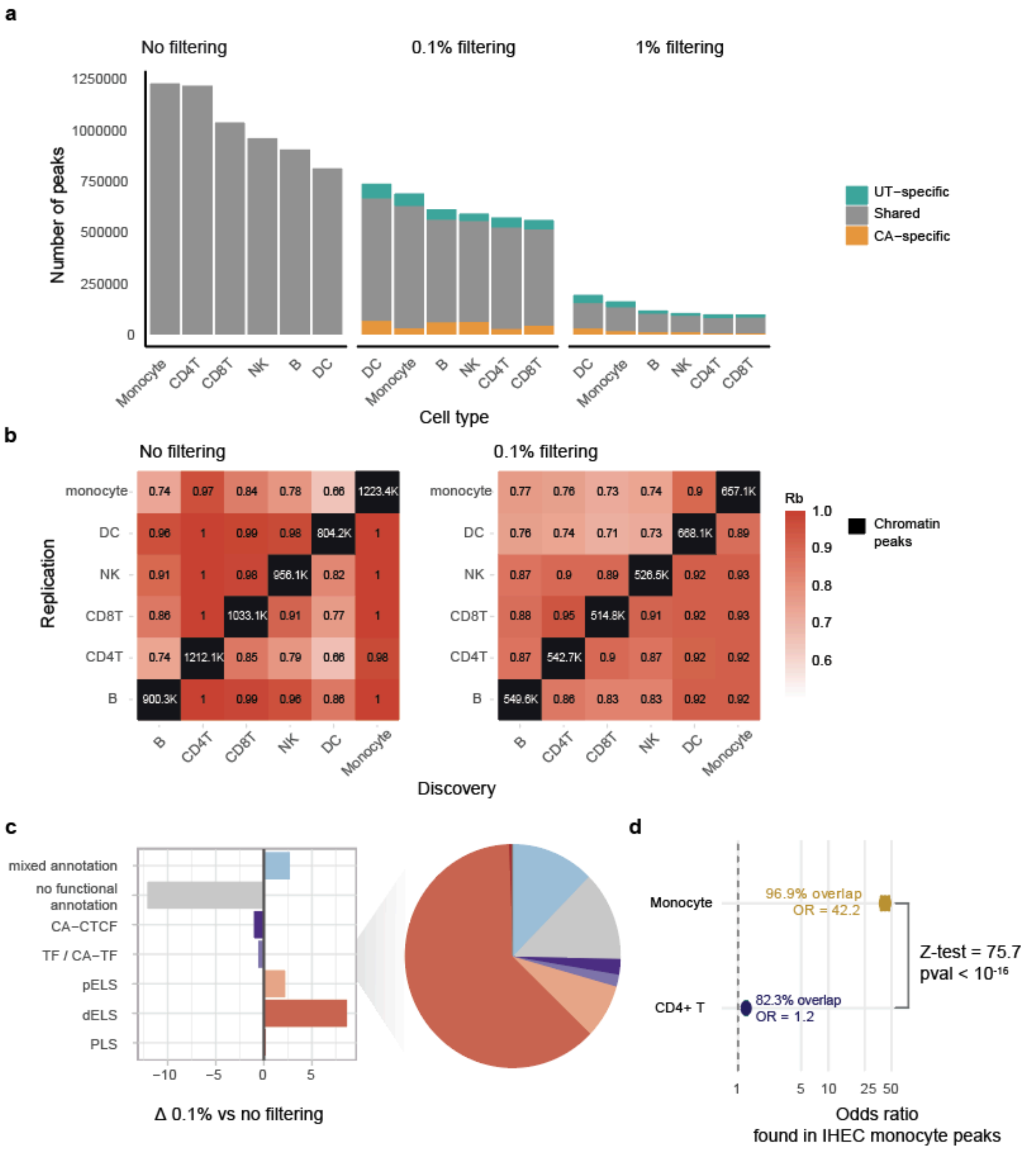


***Supplementary Note Figure 1. a.*** *Number of consensus open chromatin regions (cPeaks) per cell type that remain after filtering for peaks present in at least 0%, 0.1%, or 1% of all peaks called in that cell type.* ***b.*** *Heatmap showing the percentage of cPeaks that overlap across cell types (with no filter or a 0.1% filter applied), where the cell type in which the peak is discovered is on the x-axis, and the peak in which it is replicated is on the y-axis.* ***c.*** *ENCODE Screen v4 peak annotation (PLS: promoter-like sequence, dELS: distal enhancer-like sequence, pELS: proximal enhancer-like sequence, Other: all other categories than PLS, dELS, PLS)*[3] *of all cPeaks (without filter or a 0.1% filter applied) that are found in at least one cell type.*

### Supplementary Figures

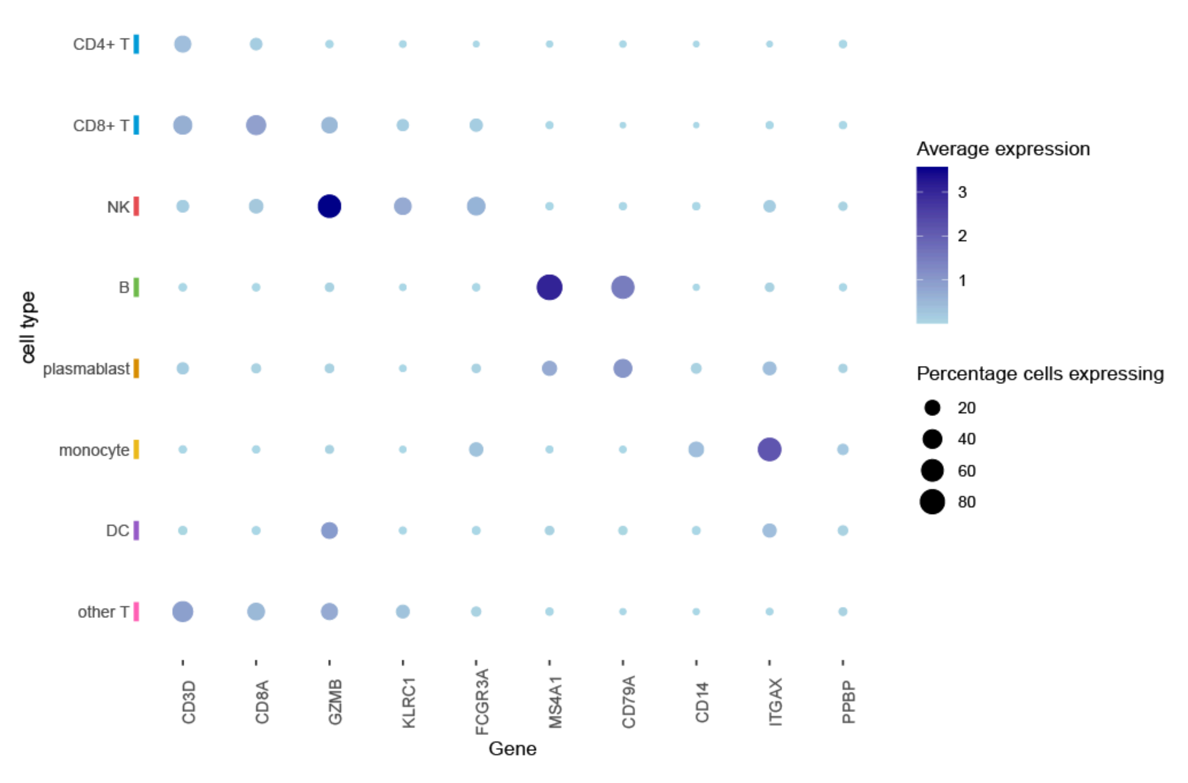


***Supplementary Figure 1.*** *Marker gene expression of cell types annotated using Azimuth[4] reference based mapping.*

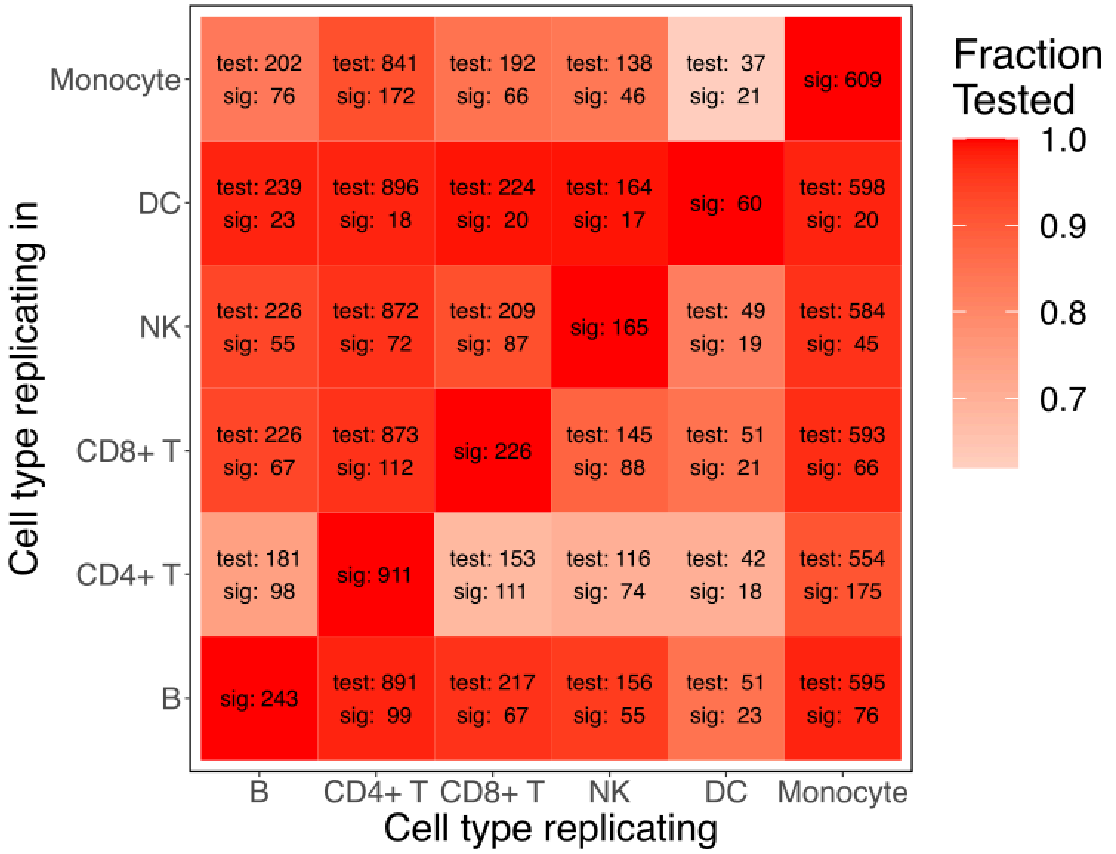


***Supplementary Figure 2.*** *Heatmap showing the number of testable and significant top-eQTLs discovered in the cell type on the x-axis and nominally-replicated in the cell type on the y-axis.*

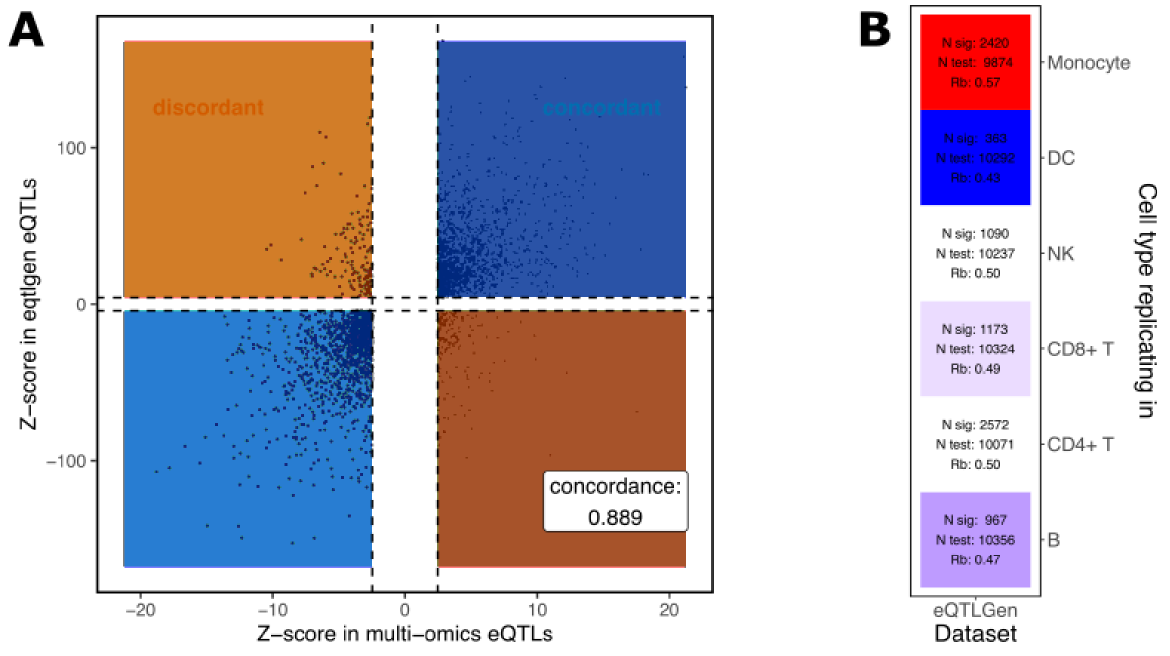


***Supplementary Figure 3. a.*** *Scatter plot showing the eQTL Z-score concordance of the eQTLs that were identified in whole blood bulk eQTLGen phase 1 and could be significantly replicated in our scMO CD4+ T cells.* ***b.*** $r_b$ *metric describing concordance of top-eQTLs identified in whole blood bulk eQTLgen phase 1*[5] *and nominally-replicated in our scMO dataset.*

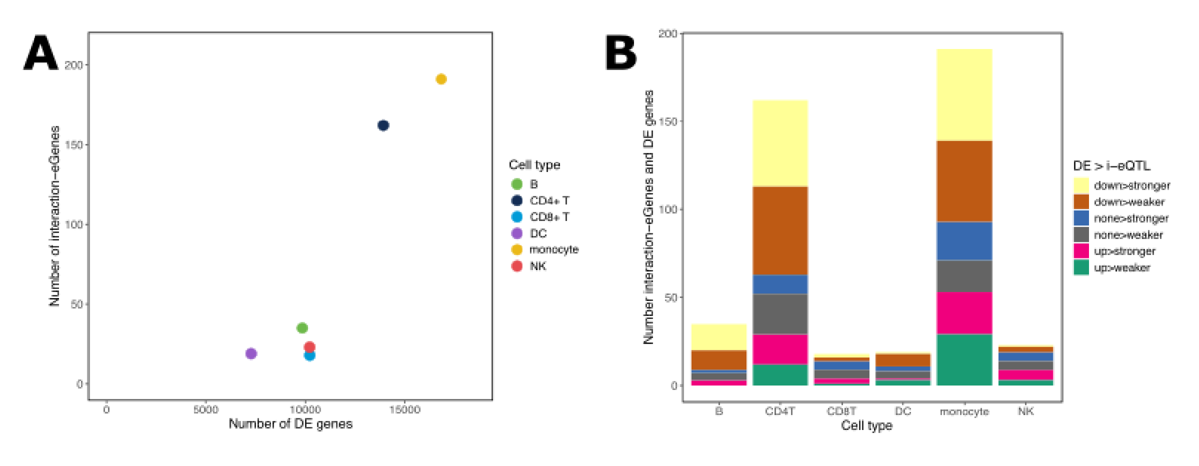


***Supplementary Figure 4. a.*** *Number of differentially expressed (DE) genes upon 24h C. albicans (24h CA) stimulation versus the number of stimulation-interaction-eGenes per cell type.* ***b.*** *Gene annotation of the top stimulation-interaction-eQTLs by the DE direction (down, none, up) and the stimulation-interaction-eQTL effect direction (stronger, weaker) by which the gene changes upon 24h CA stimulation.*

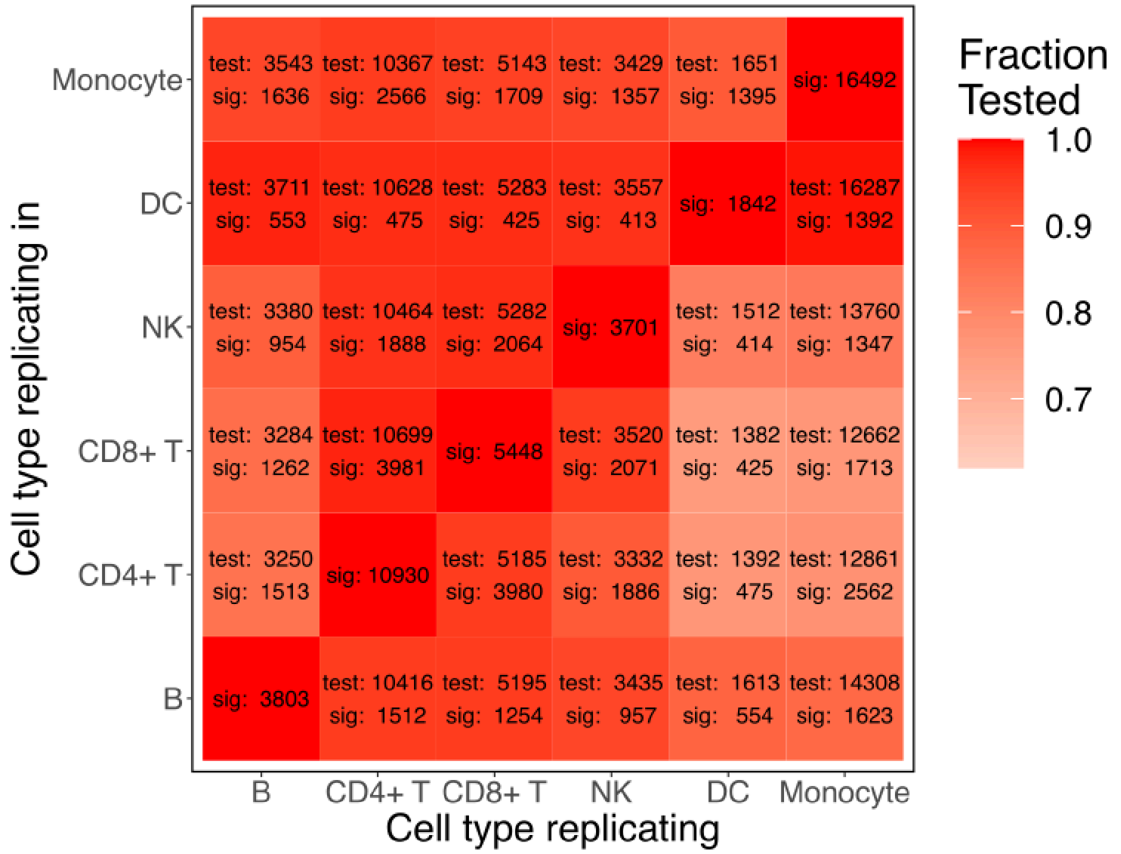


***Supplementary Figure 5.*** *Heatmap showing the number of testable and significant top-caQTLs discovered in the cell type on the x-axis and nominally-replicated in the cell type on the y-axis.*

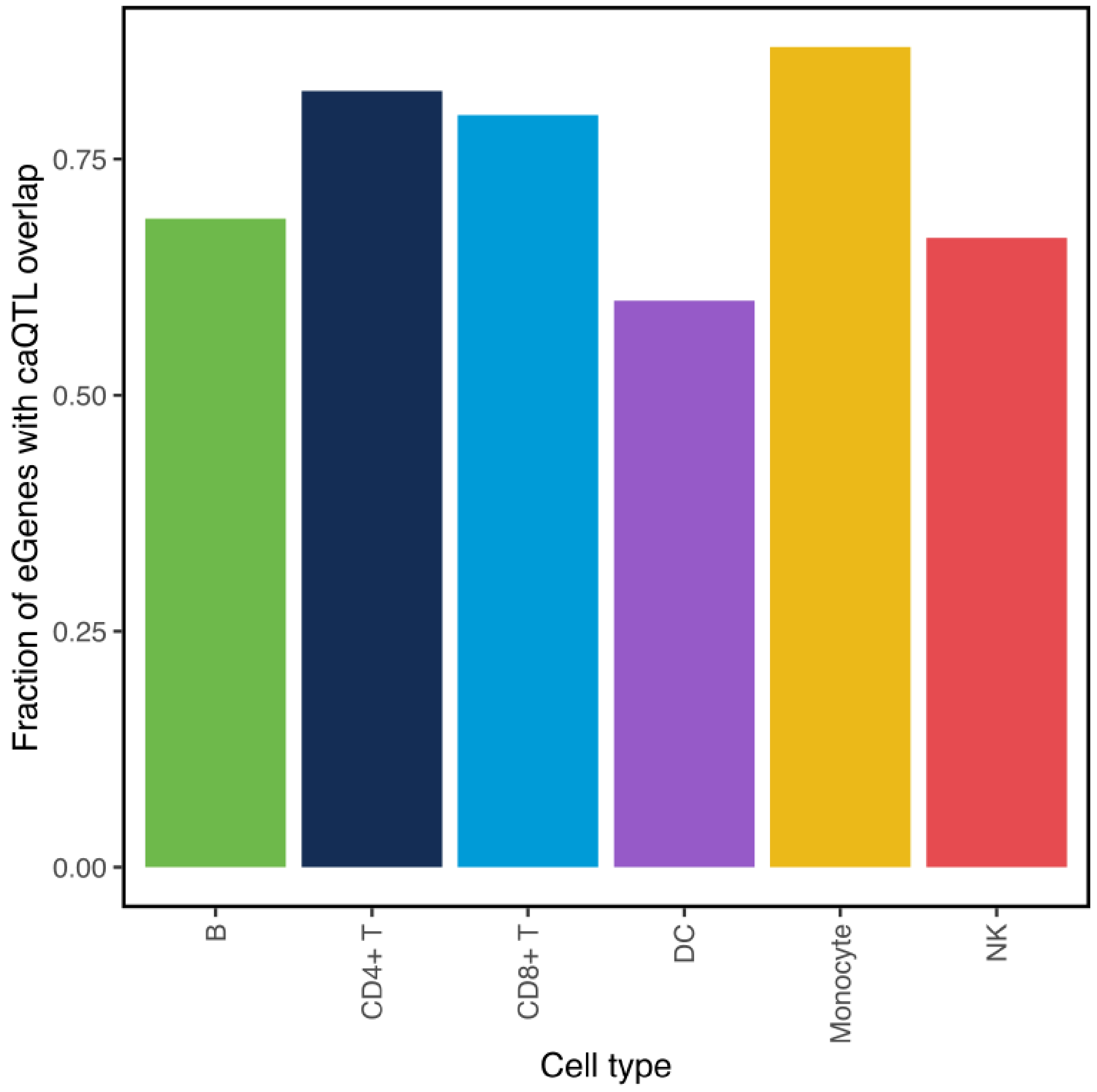


***Supplementary Figure 6.*** *Fraction of eGenes whereby the eSNP acts as dual QTL (both a caQTL and an eQTL) in the matching cell type.*

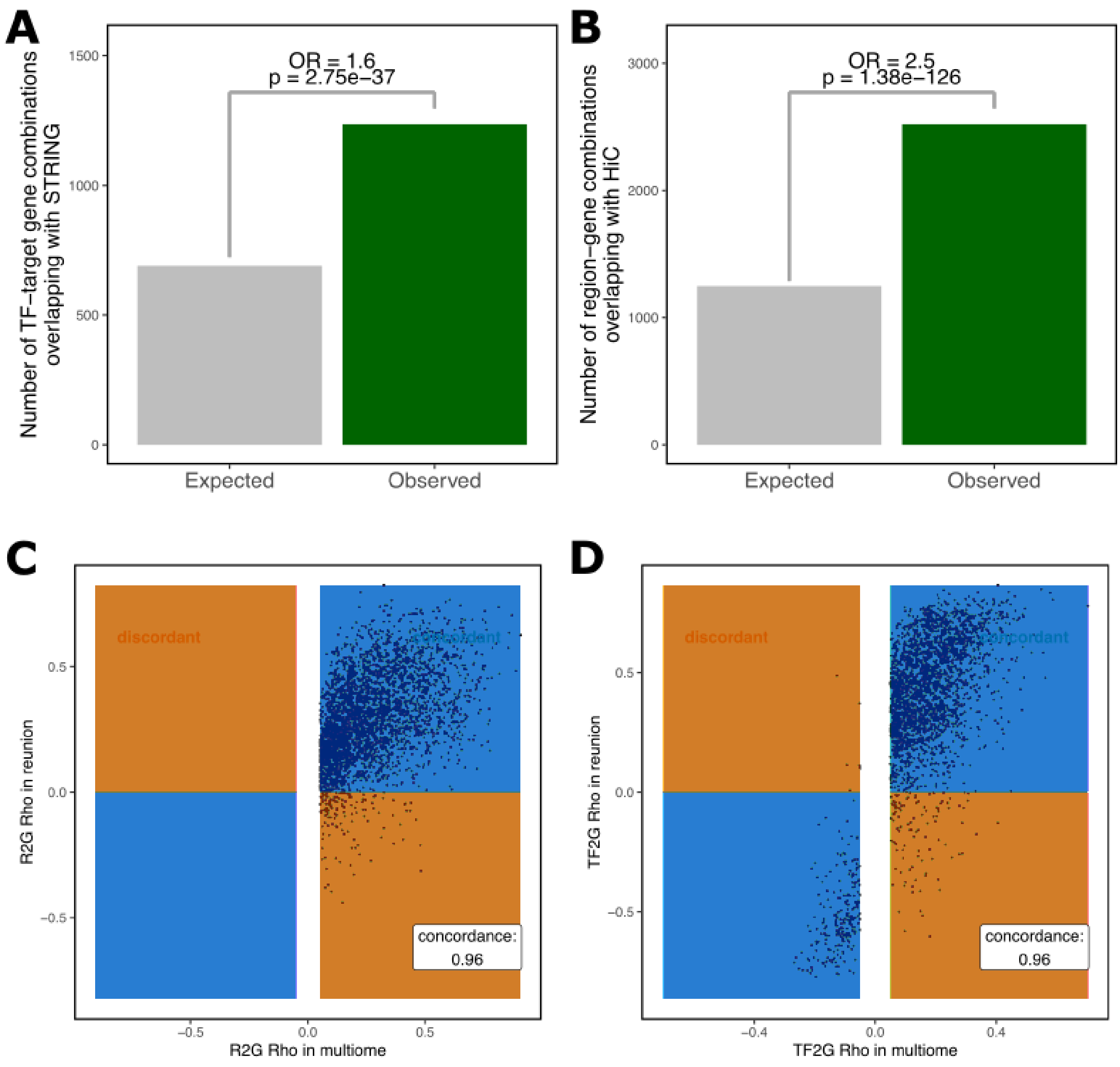


***Supplementary Figure 7. a.*** *The expected versus observed number of overlapping TF-target gene combinations between eRegulons and protein-protein interactions from the STRING database[6].* ***b.*** *The expected versus observed number of overlapping peak-gene combinations between eRegulons and K562 HiC data.* ***c.*** *Scatter plot showing the concordance of region-gene links (R2G rho) identified in our PBMC scMO SCENIC+ data versus previously published R2G rho links identified using REUNION in a PBMC sc-multiomics dataset[7].* ***d.*** *Scatter plot showing concordance of tf-gene links (TF2G rho) identified as described in c.*

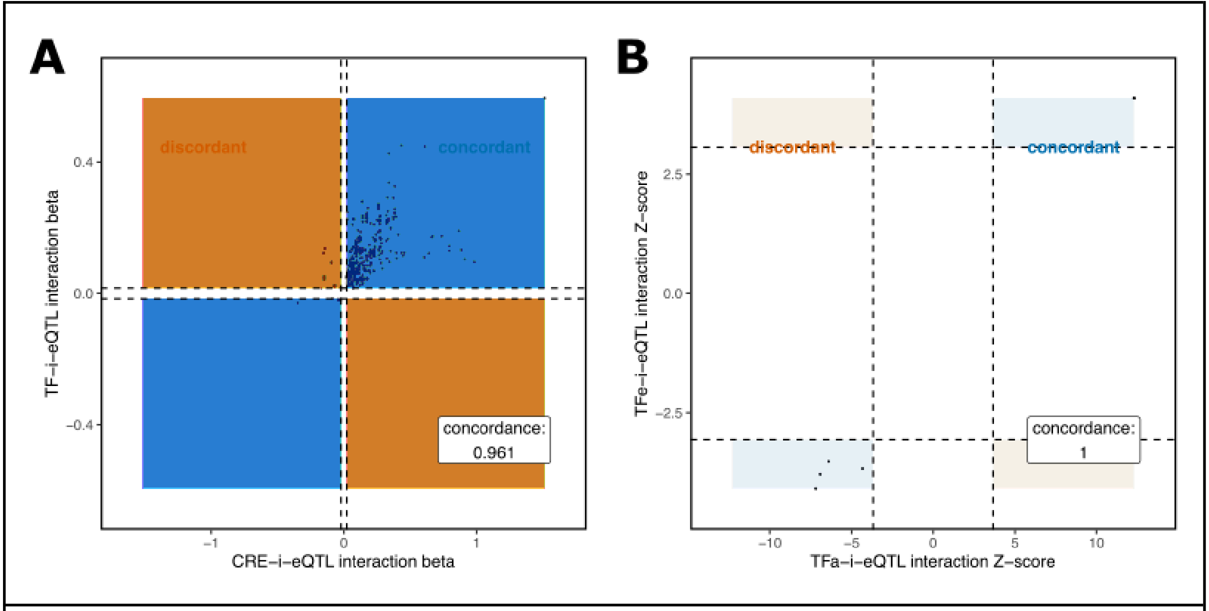


***Supplementary Figure 8. a.*** *Scatter plot showing the concordance of CRE-interaction-eQTLs with TFactivity-interaction-eQTL effect sizes.* ***b.*** *Scatter plot showing the concordance of significant TFactivity-i-eQTL effect sizes versus nominally significant TFexpression-i-eQTL effect sizes.*

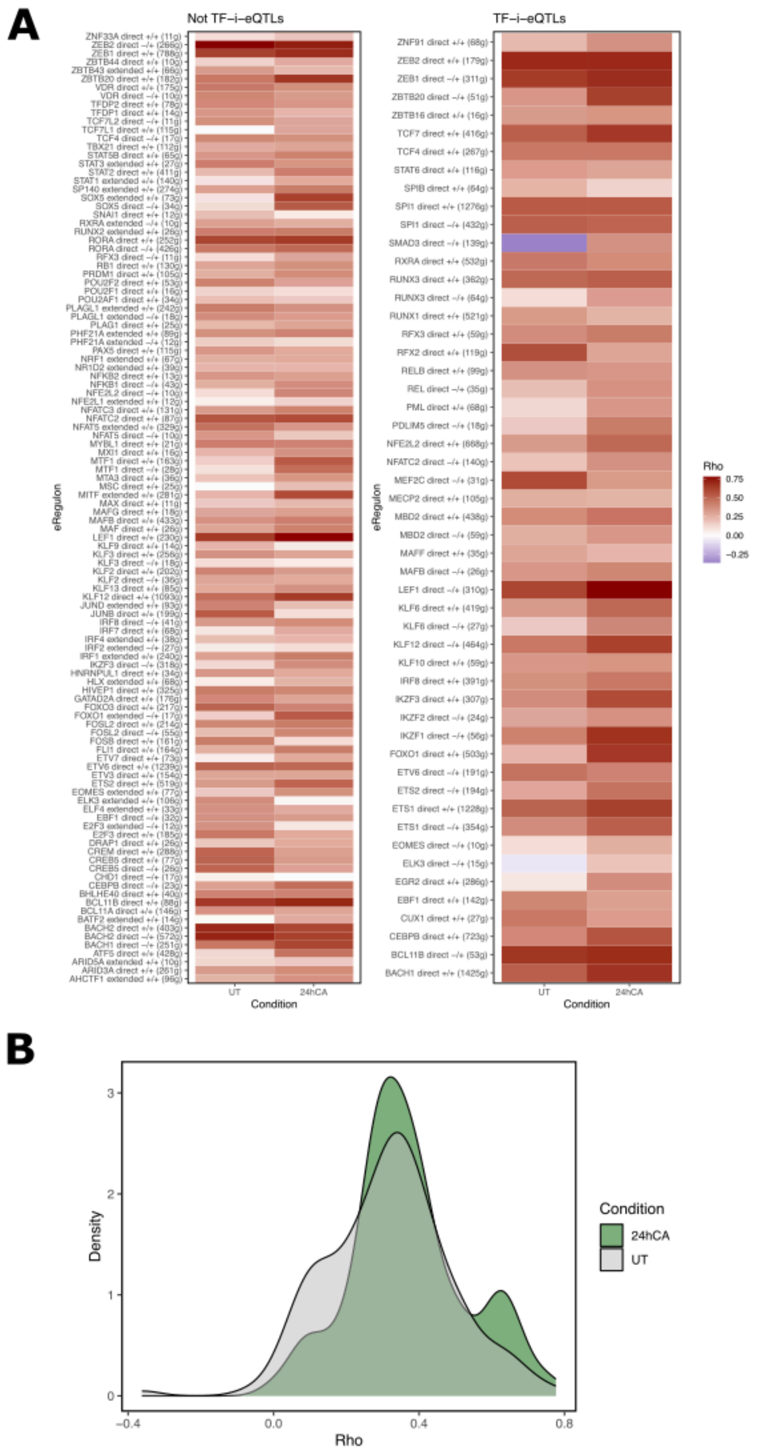


***Supplementary Figure 9***. ***a***. *Spearman correlations between gene expression and gene activity as ascertained by SCENIC+*[8]*, split by presence of TFactivity-i-eQTL status.* ***b***. *Distribution of Spearman rho correlations between TF activity and TF expression split by stimulation condition (unstimulated (UT), 24h C. albicans (24hCA)).*
